# Orai mediated Calcium entry determines activity of central dopaminergic neurons by regulation of gene expression

**DOI:** 10.1101/2023.05.24.542083

**Authors:** Rishav Mitra, Shlesha Richhariya, Gaiti Hasan

## Abstract

Maturation and fine-tuning of neural circuits frequently requires neuromodulatory signals that set the excitability threshold, neuronal connectivity and synaptic strength. Here we present a mechanistic study of how neuromodulator stimulated intracellular Ca^2+^ signals, through the store - operated Ca^2+^ channel Orai, regulate intrinsic neuronal properties by control of developmental gene expression in flight promoting central dopaminergic neurons (fpDANs). The fpDANs receive cholinergic inputs for release of dopamine at a central brain tripartite synapse that sustains flight (Sharma and Hasan, 2020). Cholinergic inputs act on the muscarinic acetylcholine receptor to stimulate intracellular Ca^2+^ release through the endoplasmic reticulum (ER) localised inositol 1,4,5-trisphosphate receptor followed by ER-store depletion and Orai mediated store-operated Ca^2+^ entry (SOCE). Analysis of gene expression in fpDANs followed by genetic, cellular and molecular studies identified Orai-mediated Ca^2+^ entry as a key regulator of excitability in fpDANs during circuit maturation. SOCE activates the transcription factor Trithorax-like (Trl) which in turn drives expression of a set of genes including *Set2*, that encodes a histone 3 Lysine 36 methyltransferase (H3K36me3). Set2 function establishes a positive feedback loop, essential for receiving neuromodulatory cholinergic inputs and sustaining SOCE. Chromatin modifying activity of Set2 changes the epigenetic status of fpDANs and drives expression of key ion channel and signaling genes that determine fpDAN activity. Loss of activity reduces the axonal arborisation of fpDANS within the MB lobe, and prevents dopamine release required for maintenance of long flight.

**Highlights:**

1. Store-operated Ca^2+^ entry (SOCE) through Orai is required in a set of flight-promoting central dopaminergic neurons (fpDANs) during late pupae and early adults to establish their gene expression profile.
2. SOCE activates a homeobox transcription factor, ‘*Trithorax-like*’ and thus regulates expression of histone modifiers Set2 and *E(z)* to generate a balance between opposing epigenetic signatures of H3K36me3 and H3K27me3 on downstream genes.
3. SOCE drives a transcriptional feedback loop to ensure expression of key genes required for neuronal function including the muscarinic acetylcholine receptor (*mAChR*) and the inositol 1,4,5-trisphosphate receptor (*itpr)*.
4. The transcriptional program downstream of SOCE is key to functional maturation of the dopaminergic neurons, enabling their neuronal excitability, axonal arborization and synaptic transmission required for adult flight.

## Introduction

Neural circuitry underlying mature adult behaviours emerges from a combination of developmental gene expression programs and experience-dependent neuronal activity. Holometabolous insects such as *Drosophila*, reconfigure their nervous system during metamorphosis to generate circuitry capable of supporting adult behaviours (Levine, 1984). During pupal development, neurons undergo maturation of their electrical properties with a gradual increase in depolarizing responses and consequent synaptic transmission (Hardie et al., 1993; Jarvilehto and Finell, 1983). Apart from voltage-gated Ca^2+^ channels and fast-acting neurotransmitters specific to ionotropic receptors, the developing nervous system also uses neuromodulators, that target metabotropic receptors, show slower response kinetics and are capable of affecting a larger subset of neurons by means of diffusion aided volumetric transmission (Taber and Hurley, 2014). Although neuromodulators alter intrinsic neuronal properties (Marder, 2012) by changes in gene expression, the molecular mechanisms through which this is achieved and maintained over developmental timescales needs further understanding.

Ca^2+^ signals generated by neuronal activity can determine neurotransmitter specification (Spitzer, 2012), synaptic plasticity (O’Hare et al., 2022; Takechi et al., 1998), patterns of neurite growth (Gu and Spitzer, 1995) and gene expression programs (Ciceri et al., 2022; Rosenberg and Spitzer, 2011) over developmental timescales (McKinney et al., 2022) where output specificity is defined by signal dynamics. Neuromodulators generate intracellular Ca^2+^ signals by stimulation of cognate G-Protein Coupled Receptors (GPCRs) linked to the inositol 1,4,5-trisphosphate (IP_3_) and Ca^2+^ signalling pathway. IP_3_ transduces intracellular Ca^2+^ release through the endoplasmic reticulum (ER) localised ligand-gated ion channel, the inositol 1,4,5-trisphosphate receptor (IP_3_R; Streb et al., 1984), followed by store-operated Ca^2+^ entry (SOCE) through the plasma membrane localised Orai channel (Prakriya and Lewis, 2015; Thillaiappan et al., 2019). Cellular and physiological consequences of Ca^2+^ signals generated through SOCE exhibit different time-scales and sub-cellular localisation from activity induced signals, suggesting that they alter neuronal properties through novel mechanisms.

Expression studies, genetics and physiological analysis support a role for IP_3_/Ca^2+^ and SOCE in neuronal development and the regulation of adult neuronal physiology across organisms (Hasan and Sharma, 2020; Mitra and Hasan, 2022; Somasundaram et al., 2014). In *Drosophila*, neuromodulatory signals of neurotransmitters and neuropeptides stimulate cognate GPCRs on specific neurons to generate IP_3_ followed by intracellular Ca^2+^ release and SOCE (Agrawal et al., 2013; Megha and Hasan, 2017; Shakiryanova et al., 2011). Genetic and cellular studies identified neuromodulatory inputs and SOCE as an essential component for maturation of neural circuits for flight (Pathak et al., 2015; Venkiteswaran and Hasan, 2009). A focus of this flight deficit lies in a group of central dopaminergic neurons (or DANs) that include the PPL1 and PPM3 DANs (Liu et al. 2012) marked by *THD’ GAL4,* alternately referred to henceforth as flight promoting DANs (fpDANs). Among the fpDANs some PPL1 DANs project to the γ2α’1 lobe of the Mushroom Body (MB; Mao and Davis 2009) a dense region of neuropil in the insect central brain, that forms a relay centre for flight (Sharma and Hasan, 2020). Axonal projections from Kenyon cells (KC) carry sensory inputs (Cervantes-Sandoval et al., 2017; Tsao et al., 2018; Yagi et al., 2016) and those from DANs carry internal state information (Mao and Davis, 2009; Riemensperger et al., 2005; Zolin et al., 2021) to functional compartments of the MB, where dopamine release modulates the KC and MB output neuron synapse (MBONs; Aso et al. 2014). The KC-DAN-MBON tripartite synapse, carries a dynamically updating representation of the motivational and behavioural state of the animal (Aso and Rubin, 2016; Berry et al., 2015; Owald et al., 2015; Waddell, 2016). Here we have investigated the molecular mechanisms that underlie the ability of neuromodulatory acetylcholine signals, acting through SOCE during circuit maturation, to determine fpDAN function required for *Drosophila* flight.

## Results

### A spatio-temporal requirement for SOCE determines flight

To understand how neuromodulator stimulated SOCE alters neuronal properties over developmental time scales we began by refining further the existing spatio-temporal coordinates of SOCE requirement for flight. Flight promoting dopaminergic neurons (DANs) with a requirement for SOCE have been identified earlier by expression of a dominant negative *Orai* transgene, which renders the channel Ca^2+^ impermeable, thus abrogating all SOCE as observed in primary neuronal cultures (*Orai^E180A^*; Figure 1A; Pathak et al. 2015; Yeromin et al. 2006). These include a smaller subset of *THD’GAL4* marked DANs, where acetylcholine promotes dopamine release through the muscarinic acetylcholine receptor (mAchR) by stimulating IP_3_/Ca^2+^ signaling (Sharma and Hasan, 2020). *Orai^E180A^* expression in a subset of 21-23 dopaminergic neurons marked by *THD’ GAL4* (Figure 1B) led to complete loss of flight (Figure 1C) while overexpression of the wildtype *Orai* transgene had a relatively minor effect (Figure 1-figure supplement 1).

**Figure 1.**
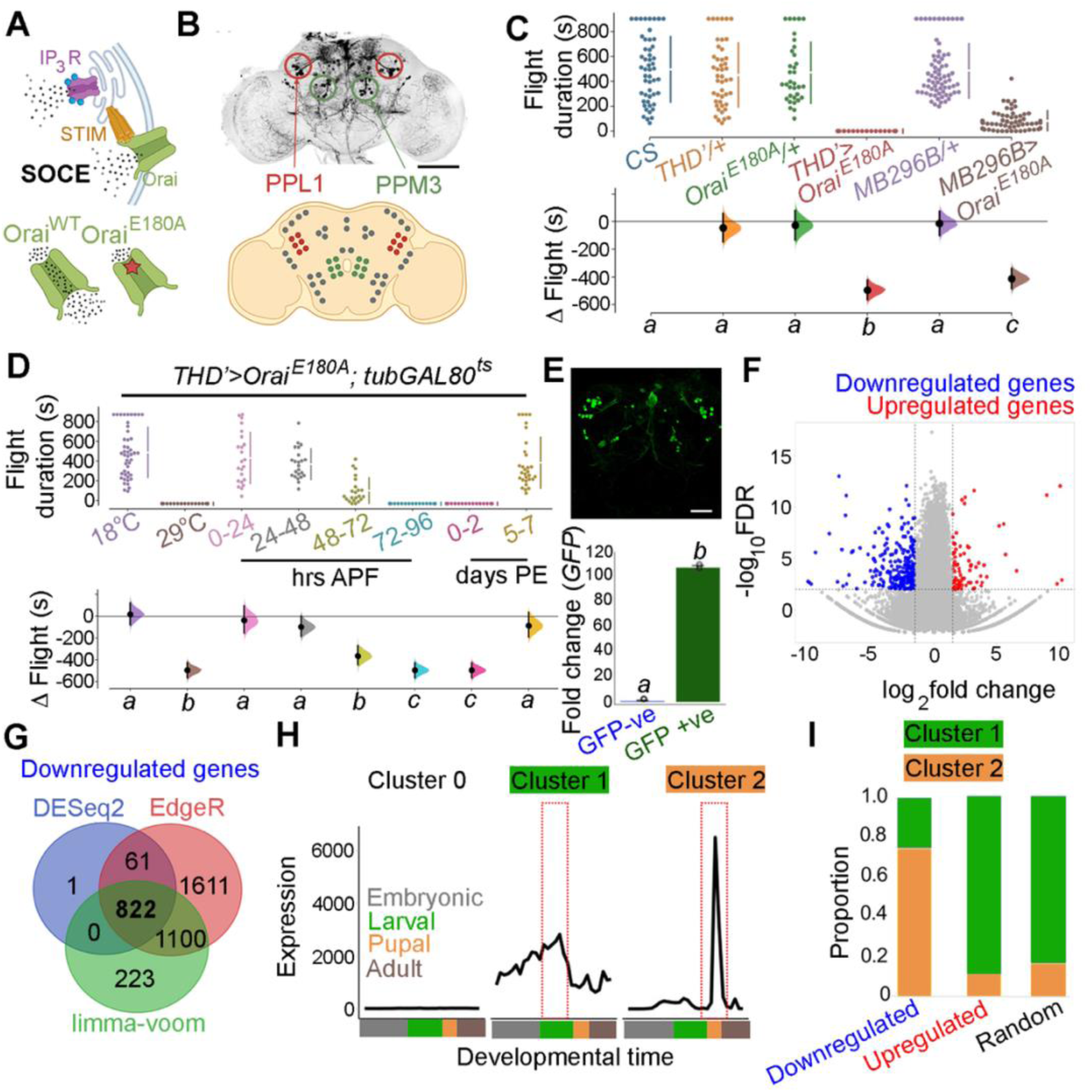
Orai-mediated Ca^2+^ entry sets the gene expression profile of flight-promoting dopaminergic neurons in late development and early adulthood. **A**. A schematic of Ca^2+^ release through the IP_3_R and SOCE through STIM/Orai (upper panel) followed by representation of the wildtype (Ca^2+^ permeable) and mutant (Ca^2+^ impermeable) Orai channels (lower panel). **B**. Anatomical location of *THD’* DANs in the fly central brain immunolabelled for mCD8GFP (upper panel) followed by a cartoon of central brain DAN clusters. Scale bar indicates 20 μm. PPL1 and PPM3 clusters are labelled in red and green respectively. **C**. Measurement of flight bout durations demonstrate a requirement for Orai-mediated Ca^2+^ entry in *THD’* DANs and in 2 pairs of PPL1 DANs marked by the MB296B driver. **D**. *THD*’ DANs require Ca^2+^ entry through Orai at 72-96 hours APF and 0-2 days post eclosion to promote flight. In **C** and **D** flight bout durations in seconds (s) are represented as a swarm plot where each genotype is represented by a different color, and each fly as a single data point. The Δ Flight parameter shown below indicates the mean difference for comparisons against the shared Canton S control and is shown as a Cumming estimation plot. Mean differences are plotted as bootstrap sampling distributions. Each 95% confidence interval is indicated by the ends of the vertical error bars. The same letter beneath each distribution refers to statistically indistinguishable groups after performing a Kruskal-Wallis test followed by a post hoc Mann-Whitney U-test (p<0.005). At least 30 flies were tested for each genotype. **E**. *THD’* DANs were labelled with cytosolic eGFP (10 μm scale), isolated using FACS and validated for enrichment of *GFP* mRNA by qRT-pCR (lower panel). The qRT-pCR results are from 4 biological replicates with different letters representing statistically distinguishable groups after performing a two-tailed t test (p<0.05). **F.** RNA-seq comparison of FAC sorted populations of GFP labelled THD’ DANs from *THD’>GFP* and *THD’>GFP;Orai^E180A^* pupal dissected CNSs. The RNA-Seq data is represented in the form of a volcano plot of fold change vs FDR. Individual dots represent genes, coloured in red (upregulated) or blue (downregulated) by >1 fold. **G.** Downregulated genes were identified by 3 different methods of DEG analysis, quantified and compared as a Venn diagram. **H.** Gene expression trajectories of SOCE-induced DEGs plotted as a function of developmental time (modENCODE Consortium et al., 2010) and clustered into 3 groups using k-means analysis. **I**. The relative proportion of downregulated, upregulated genes, and a random set of genes found in the clusters described in (**H**) indicate that 75% of downregulated genes exhibit a pupal peak of expression.

*THD’* marks DANs in the PPL1, PPL2, and PPM3 clusters of the adult central brain (Figure 1B) of which the PPL1 (10-12) and PPM3 (6-7) cell clusters constitute the fpDANs (Pathak et al., 2015). Among these 16-19 DANs, two pairs of PPL1 DANs projecting to the γ2α’1 lobes of the mushroom body (marked by the MB296B split GAL4 driver and schematised in Figure 1-figure supplement 2; Aso et al. 2014; Aso and Rubin 2016) were identified as contributing to the flight deficit to a significant extent (Figures 1C; Figure 1-figure supplement 3). The MB296B DANs form part of a central flight relay centre identified earlier as requiring IP_3_/Ca^2+^ signaling (Sharma and Hasan, 2020). That the *THD’* neurons are required for flight was further confirmed by inhibiting their function. Acute activation of either a temperature sensitive *Dynamin* mutant (*Shibire^ts^*; Kitamoto 2001), which blocks exocytosis at an elevated temperature, or acute expression of the tetanus toxin light chain fragment (*TeTxLC*; Sweeney et al. 1995) which cleaves *Synaptobrevin*, an essential component of the synaptic release machinery in *THD’* neurons led to severe flight defects (Figure 1-figure supplement 4).

Previous work has demonstrated that intracellular Ca^2+^ release through the IP_3_R, that precedes SOCE (Figure 1A), is required during late pupal development in *THD’* marked DANs for flight (Pathak et al., 2015; Sharma and Hasan, 2020). The precise temporal requirement for SOCE in *THD’* neurons was investigated by inducing *Orai^E180A^* transgene expression for specific periods of pupal development and in adults with the *tubulinGAL80^ts^* based temperature sensitive expression system, (TARGET-Temporal And Regional Gene Expression Targeting; McGuire et al., 2004). Approximately 100 hours of pupal development were binned into 24 hour windows, wherein SOCE was abrogated by *Orai^E180A^* expression, following which, normal development and growth was permitted. Flies were assayed for flight 5 days post eclosion. Abrogation of SOCE by expression of the *Orai^E180A^* dominant negative transgene at 72 to 96 hrs after puparium formation (APF), resulted in complete loss of flight as compared to minor flight deficits observed upon expression of *Orai^E180A^* during earlier developmental windows (Figure 1D). Abrogation of flight also occurred upon *Orai^E180A^* expression during the first 2 days post-eclosion (see discussion) whereas abrogation of SOCE 5 days after eclosion as adults resulted in only modest flight deficits (Figure 1D), indicating that SOCE is required for maturation of *THD’* marked flight dopaminergic neurons. Earlier work has shown that loss of SOCE by expression of *Orai^E180A^* does not alter the number of dopaminergic neurons or affect neurite projection patterns of PPL1 and PPM3 neurons (Pathak et al., 2015). Of these two clusters, PPM3 DANs project to the ellipsoid body in the central complex (Kong et al., 2010), and PPL1 DANs project to the mushroom body and the lateral horn (Mao and Davis, 2009) and aid in the maintenance of extended flight bouts (Sharma and Hasan, 2020).

### Loss of SOCE in late pupae leads to a re-organisation of neuronal gene expression

The molecular consequences of loss of SOCE in *THD’* neurons were investigated next by undertaking a comprehensive transcriptomic analysis from fluorescence activated and sorted (FACS) *THD’* neurons with or without *Orai^E180A^* expression and marked with eGFP (Figure 1E; Figure 1-figure supplement 5). *THD’* neurons were obtained from pupae at 72 ± 6h APF (essential SOCE requirement for flight; Figure 1E). Expression analysis of RNAseq libraries generated from *THD’* neurons revealed a reorganisation of the transcriptome upon loss of SOCE (Figure 1F). Expression of 822 genes was downregulated (with a fold change cut-off of < −1) as assessed using 3 different methods of Differentially Expressed Genes (DEG) analysis (Figure 1G) whereas 137 genes were upregulated (Figure 1-figure supplement 6). To understand if loss of SOCE affects genes expressed in neurons from the pupal stage, we used the modEncode dataset (Celniker et al., 2009; modENCODE Consortium et al., 2010) to reconstruct developmental trajectories of all genes expressed in *THD’* neurons (Figure 1-figure supplement 7). Using an unsupervised clustering algorithm, we classified genes expressed in *THD’* neurons into 3 clusters where ‘Cluster 0’ is low expression throughout development; ‘Cluster 1’ exhibits a larval peak in expression and ‘Cluster 2’ exhibits a pupal peak in expression (Figure 1H). The majority of expressed genes (>95%) classify as low expression throughout (Cluster 0) and were removed from further analysis. Genes belonging to either the downregulated gene set or the upregulated gene set in SOCE-deficient *THD’* neurons and a random set of genes were analysed further. A comparison of the proportion of DEGs classified in the larval and pupal clusters revealed that >75% of downregulated genes belonged to the pupal peak (Cluster 2) whereas <20% of either the upregulated genes or a random set of genes classified as belonging to the larval peak (Figure 1I). Moreover, downregulated genes exhibit higher baseline expression in wildtype pupal *THD’* neurons (Figure 1-figure supplements 8-9). These analyses suggest that SOCE induces expression of a set of genes in *THD’* neurons during the late pupal phase which are subsequently required for flight. These are henceforth referred to as SOCE-responsive genes.

### SOCE regulates gene expression through a balance of Histone 3 Lysine 36 trimethylation and Histone 3 Lysine 27 trimethylation

Gene ontology (GO) analysis of SOCE-responsive genes revealed ‘Transcription’, ‘Ion transporters’, ‘Ca^2+^ dependent exocytosis’, ‘GPCRs’, ‘Synaptic components’, and ‘Kinases’ as top GO categories (Figure 2A). Among these, ‘Transcription’ and ‘Ion transporters’ represented categories with the highest enrichment. Further analysis of genes that classified under ‘Transcription’ revealed SET domain containing genes that encode histone lysine methyltransferases implicated in chromatin regulation and gene expression (Dillon et al., 2005) and among the SET domain containing genes, *Set1* and *Set2* showed distinct downregulation (Figure 2B). One of these genes, the H3K36me3 methyltransferase *Set2* (Figure 2C), was identified in a previous RNAseq from larval neurons with loss of IP_3_R-mediated Ca^2+^ signaling (Mitra et al., 2020). RT-PCRs from sorted *THD’* neurons confirmed that *Set2* levels are indeed downregulated (by 76%) in *THD’* neurons expressing *Orai^E180A^* and to the same extent as seen upon expression of *Set2^RNAi^* (79% downregulation; Figure 2C). *Set1* and *Set2* perform methyltransferase activity at H3K4 and H3K36 histone residues respectively (Ardehali et al., 2011; Stabell et al., 2007). Both these markers are epigenetic signatures for transcriptional activation, where H3K4me3 is enriched at the 5’ end of the gene body and H3K36me3 is enriched in the gene bodies of actively transcribed genes; schematised in Figure 2-figure supplement 1). In the adult CNS, *Set2*, is expressed at 4-fold higher levels compared to *Set1* (Figure 2-figure supplement 2; modENCODE Consortium et al., 2010). *Set2* also shows a steep 5.5 fold increase in expression levels during the pupal to adult transition, compared to a 2 fold increase in *Set1* (Figure 2-figure supplement 2). These observations coupled with the previous report of Set2 function downstream of IP_3_R/Ca^2+^ signaling in larval glutamatergic neurons (Mitra et al., 2020), led us to pursue the role of Set2 in *THD’* neurons.

**Figure 2.**
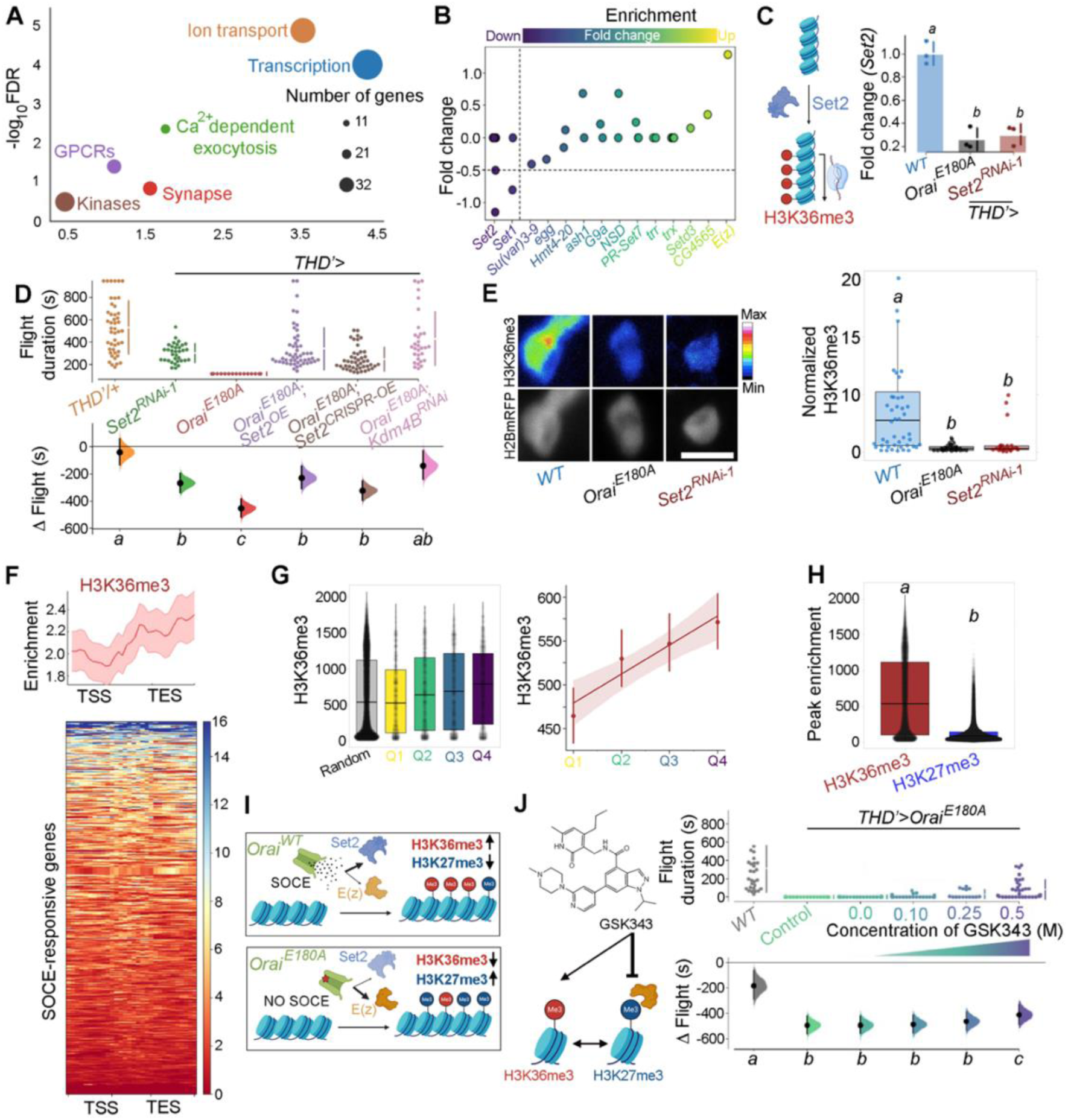
Ca^2+^ entry through Orai regulates gene expression by Set2-mediated histone modification. **A**. Scatterplot of GO categories enriched in SOCE-responsive genes. Individual GO terms are represented as differently colored circles, with radius size indicating number of genes enriched in that category. **B**. Fold change of SET domain containing genes as indicated. Individual circles on the Y-axis for each gene represent transcript variants pertaining to that gene. **C**. Transcripts of the H3K36 methyltransferase (left) are significantly diminished (right) in *THD’* DANs either upon loss of Orai function (*THD’>Orai^E180A^*) or by knockdown of *Set2* (*THD’>Set2^RNAi^*). qRT-PCRs were performed from FAC sorted *THD’* DANs with 3 biological replicates. Individual 2^-ΔΔCT^ values are shown as points. Letters represent statistically distinguishable groups after performing an ANOVA and post hoc Tukey test (p<0.05). **D**. Significant rescue of flight bout durations seen in *THD’ > Orai^E180A^* flies by overexpression of *Set2* and by knockdown of the *Kdm4B* demethylase indicating a net requirement for H3K36me3. Flight durations of single flies are depicted as swarm plots and the Δ flight parameter is shown below. Both were measured as described in the legend to Figure 1. N = 30 or more flies for each genotype. Letters represent statistically distinguishable groups after performing an ANOVA and post hoc Tukey test (p<0.005) **E**. Representative images (upper panel) and quantification (lower panel) from immunostaining of H3K36me3 and H2BmRFP in nuclei of *THD’* DANs from at least 10 brains. Scale bar represents 5 μm. The boxplot represents individual H3K36me3/H2BmRFP ratios from each *THD’* DAN for each genotype. Letters represent statistically distinguishable groups after performing an ANOVA and post hoc Tukey test (p<0.05) **F**. H3K36me3 enrichment over the gene bodies of SOCE-responsive genes in wildtype fly heads represented in the form of a tag density plot. **G**. H3K36me3 signal is enriched on WT SOCE-responsive genes with greater downregulation upon loss of Orai function. Individual data points represent *WT* H3K36me3 ChIP-Seq signals from adult fly heads represented as a boxplot (left) and a regression plot (right; Pearson’s correlation coefficient = 0.11) indicating a correlation between extent of downregulation upon loss of SOCE and greater enrichment of H3K36me3. **H**. SOCE-responsive genes are enriched in H3K36me3 signal as compared to H3K27me3 signal as measured from relevant ChIP-Seq datasets. Adult fly head ChIP-Seq datasets for measurements in **F** to **H** were obtained from modEncode (modENCODE Consortium et al., 2010). **I**. Schematic representation of how Orai mediated Ca^2+^ entry regulates a balance of two opposing epigenetic signatures in developing DANs. **J**. Pharmacological inhibition of H3K27me3 using GSKS343 (left) in *THD’>Orai^E180A^* flies results in a dose-dependent rescue of flight bout durations (right). Flight assay measurements from N> 30 flies, are represented as described earlier. Letters represent statistically distinguishable groups after performing an ANOVA and post hoc Tukey test (p<0.005).

*Set2* encodes the only *Drosophila* methyltransferase which is specific for H3K36 trimethylation (Stabell et al., 2007). To understand the functional significance, if any, of down-regulation of *Set2* in *THD’* neurons upon loss of SOCE, four independent *Set2* RNAi constructs were obtained. Knockdown of *Set2* in *THD’* neurons with all four RNAi lines tested resulted in significant flight defects (Figure 2D; Figure 2-figure supplement 3). *Set2* function for flight was additionally verified in the PPL1 DANs projecting to the γ2α’1 lobe of the mushroom body (*MB296BGAL4*; Figure 2-figure supplement 4). To test the hypothesis that *Set2* expression and function for flight is regulated by SOCE, we overexpressed *Set2* in the *Orai^E180A^* background, using GAL4-UAS driven heterologous *Set2^WT^* expression or CRISPR-dCas9::VPR driven overexpression (Gilbert et al., 2013) from the endogenous locus, and measured flight (Figure 2D data in purple and brown). Flies expressing *Orai^E180A^* in the *THD’* neurons, which are flightless, exhibit significant rescue in flight bout durations upon *Set2* overexpression, through either method (Figure 2D, compare red with purple and brown data), but not upon expression of a control transgene, *GCamP6m* (Figure 2-figure supplement 5, compare data in green and red). Although the *Set2* overexpression rescues were induced all through development, because Orai function drives critical gene expression during 72-96 hrs APF (Figure 1D), we concluded that Set2-mediated rescue is also likely to occur during this time window. These data taken together with downregulation of *Set2* upon expression of *Orai^E180A^* (Figure 2C) support the hypothesis that SOCE driven expression of *Set2* in fpDANs is required for flight. That Set2 function downstream of SOCE, occurs specifically through its methyltransferase activity was supported through experiments where knockdown of the demethylase *Kdm4B*, the primary histone demethylase expressed in these neurons (Figure 2-figure supplement 6), rescued flight in *THD’>Orai^E180A^* flies (Figure 2D; data in pink). Knockdown of the lower expressed *Drosophila* H3K36 demethylase isoform, *Kdm4A,* did not show a similar rescue (Figure 2-figure supplement 7; data in pink). Moreover, both *Set2* depletion and loss of SOCE (*THD’>Orai^E180A^*) lead to a decrease in the normalised H3K36me3 signal in *THD’* neurons (Figure 2E).

To understand if deficient H3K36me3 is found in the SOCE-responsive genes, we quantified the enrichment of H3K36me3 at SOCE-responsive loci from an H3K36me3 ChIP-Seq dataset (modENCODE Consortium et al., 2010) from wild-type adult heads (Figure 2F). We observed an enrichment of the H3K36me3 signal over the gene bodies of the SOCE-responsive genes (Figure 2F). Additionally, genes that were more affected by loss of SOCE (greater extent of downregulation), had a correspondingly higher H3K36me3 signal (Figure 2G) as compared with a random set of genes (Figure 2G; data in grey), further indicating that one of the ways SOCE regulates gene expression is through Set2 mediated deposition of H3K36me3.

The gene *E(z*) that encodes a component of the PRC2 complex (*EZH2*), which deposits H3K27me3 is up-regulated upon loss of SOCE (Figure 2B) suggesting that SOCE could affect additional histone modifications. While H3K36me3 is a marker for transcriptional activation (Krogan et al., 2003), H3K27me3 is a repressive signature which silences transcription (Cai et al., 2021). These two marks are antagonistic as deduced from studies which have shown that deposition of H3K36me3 allosterically inhibits the Polycomb Repressive Complex (PRC2), thereby preventing it from depositing the H3K27me3 signature at the same genomic loci (Finogenova et al., 2020). A comparison of H3K36me3 and H3K27me3 peaks over the 822 SOCE-responsive genes revealed a higher enrichment of H3K36me3 versus H3K27me3 in wild type fly heads (Figure 2H; modENCODE Consortium et al., 2010), suggesting that robust expression of these genes in adult neurons occurs through a SOCE-dependent mechanism initiated in late pupae (schematised in Figure 2I). To test this hypothesis, we fed SOCE-deficient flies (*THD’GAL4>Orai^E180A^*) GSK343, a pharmacological inhibitor (Verma et al., 2012) of the EZH2 component of PRC2 that is known to reduce H3K27me3 chromatin marks and performed flight assays. Feeding of GSK343 lead to partial rescue of flight in *THD’> Orai^E180A^* flies in a dose dependent manner with maximal flight bout durations of up to 400 sec (Figure 2J; Figure 2-figure supplement 8).

### Normal cellular responses of PPL1 DANs require Orai mediated SOCE and H3K36 trimethylation

Next we investigated Ca^2+^ responses in SOCE and H3K36 trimethylation deficient *THD’* PPL1 DANs upon stimulation with a neuromodulatory signal for the muscarinic acetylcholine receptor (mAChR). It is known that *THD’* marked PPL1 DANs receive cholinergic inputs that stimulate store Ca^2+^ release through the IP_3_R, leading to SOCE through Orai (Figure 3A; Ebihara et al., 2006; Sharma and Hasan, 2020). In support of this idea, either expression of *Orai^E180A^* or knockdown of the IP_3_R attenuated the composite ER-Ca^2+^ release plus SOCE response to carbachol (CCh), a mAChR agonist, in PPL1 DANs (Figure 3B-D; Figure 3-figure supplement 1). Ex-vivo brain preparations used here for in situ Ca^2+^ imaging of fpDANs are not viable in the absence of extracellular Ca^2+.^ Hence measurement of store-Ca^2+^ release and SOCE independent of each other was not possible. Importantly, knockdown of *Set2* also abrogated the Ca^2+^ response to CCh (Figure 3B-D), whereas overexpression of a transgene encoding *Set2* in *THD’* neurons either with loss of SOCE (*Orai^E180A^*) or with knockdown of the IP_3_R (*itpr^RNAi^*), lead to significant rescue of the Ca^2+^ response, indicating that *Set2* is required downstream of SOCE (Figure 3B-D) and IP_3_/Ca^2+^ signalling (Figure 3-figure supplement 1).

**Figure 3.**
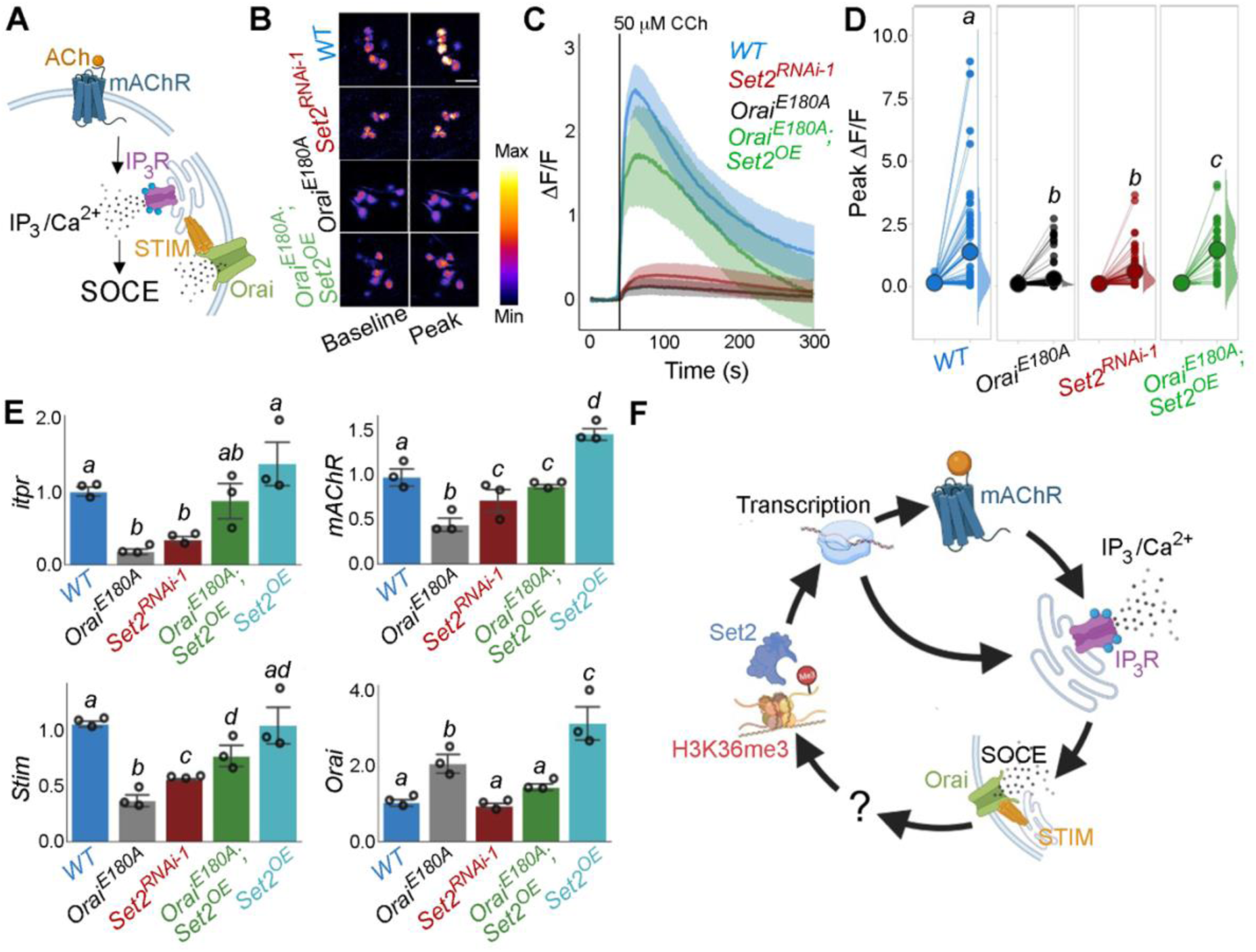
Orai-mediated Ca^2+^ entry potentiates cellular Ca^2+^ responses to cholinergic inputs through Set2 and a transcriptional feedback loop. **A**. A schematic of intracellular Ca^2+^ signaling downstream of neuromodulatory signaling where activation of mAChR stimulates intracellular Ca^2+^ release through the IP_3_R followed by SOCE through STIM/Orai. **B.** Cholinergic inputs by addition of carbachol (CCh) evoke Ca^2+^ signals as measured by change in the fluorescence of GCaMP6m in *THD’* DANs of ex-vivo brains. Representative GCaMP6m images of *THD’* DANs are shown with baseline and peak evoked responses in the indicated genotypes. Scale bar = 10 μM. **C**. Median GCaMP6m responses plotted as a function of time. A shaded region around the solid line represents the 95% confidence interval from 4-5 cells imaged per brain from 10 or more brains per genotype. **D.** Individual cellular responses depicted as a paired plot where different letters above indicate statistically distinguishable groups after performing a Kruskal Wallis Test and a Mann-Whitney U-test (p<0.05). **E**. qRT-PCR measurements of *itpr*, *mAChR*, *Stim* and *Orai* from FAC sorted *THD’* DANs obtained from three biological replicates of appropriate genetic backgrounds. The genotypes include DANs with loss of cellular Ca^2+^ responses (*THD’>Orai^E180A^*; and *THD’>Set2^RNAi-1^*), rescue of cellular Ca^2+^ response by overexpression of *Set2* (*THD’>Orai^E180A^*; *Set2^OE^*) and *Set2* overexpression. Bar plots indicate mean expression levels in comparison to *rp49*, with individual data points represented as hollow circles. The letters above indicate statistically indistinguishable groups after performing a Kruskal Wallis Test and a Mann-Whitney U-test (p<0.05). **F**. Schematic representation of a transcriptional feedback loop downstream of cholinergic stimulation in *THD’* DANs.

An understanding of how Set2 might rescue cellular IP_3_/Ca^2+^ signaling was obtained in an earlier study where it was demonstrated that *Set2* participates in a transcriptional feedback loop to control the expression of key upstream components such as the *mAChR* and the *IP_3_R* in a set of larval glutamatergic neurons (Mitra et al., 2021). To test whether *Set2* acts through a similar transcriptional feedback loop in the current context, *THD’* DANs were isolated using FACS from appropriate genetic backgrounds and tested for expression of key genes required for IP_3_/Ca^2+^ signaling (*mAChR* and *itpr)* and SOCE (*Stim* and *Orai*). Loss of SOCE upon expression of *Orai^E180A^* lead to a significant decrease in the expression of *itpr (72%* downregulation; Figure 3E) and *mAChR (56%* downregulation; Figure 3E in *THD’* DANs. As expected *Orai* expression (Figure 3E) increased 2-fold presumably due to overexpression of the *Orai^E180A^* transgene. Importantly, knockdown of *Set2* in *THD’* neurons also led to downregulation of *itpr* (43%), *mAChR* (78%) and *Stim* (61%) (Figure 3E). Overexpression of *Set2* in the background of *Orai^E180A^* led to a rescue in the levels of *itpr* (69% increase), *mAChR* (43% increase), and *Stim* (40% increase) (Figure 3E). Overexpression of *Set2* in wildtype *THD’* neurons resulted in upregulation of *mAChR* and *Orai* (Figure 3E). These findings indicate that while SOCE regulates *Set2* expression (Figure 2C), *Set2* in turn functions in a feedback loop to regulate expression of key components of IP_3_/Ca^2+^ (*mAchR* and *IP_3_R*) and SOCE *(STIM* and *Orai)* (Figure 3E). Thus, ectopic *Set2* overexpression in SOCE deficient *THD’* neurons, leads to increase in expression of key genes that facilitate intracellular Ca^2+^ signalling and SOCE (schematised in Figure 3F). Direct measures of Orai-channel function under conditions of altered *Set2* expression are needed in future to assess how the feed-back loop alters carbachol-induced ER-Ca^2+^ release and SOCE independent of each other (see ‘Limitations of this study’ in discussion). While we assume that the observed changes in gene expression translate to alterations in protein levels, direct measurement of protein levels specifically from *THD’* neurons need to be addressed in future.

### Trithorax-like or Trl is an SOCE responsive transcription factor in *THD’* DANs

The data so far identify SOCE as a key developmental regulator of the neuronal transcriptome in *THD’* marked DANs, where it upregulates *Set2* expression and thus enhances the activation mark of H3K36 trimethylation on specific chromatin regions. However, the mechanism by which SOCE regulates expression of *Set2,* and other relevant effector genes remained unresolved (Figure 3F). In mammalian T cells, SOCE leads to de-phosphorylation of the NFAT (Nuclear Factor of Activated T-cells) family of transcription factors by Ca^2+^/Calmodulin sensitive phosphatase Calcineurin, followed by their nuclear translocation and ultimately transcription of relevant target genes (Hogan et al., 2003). Unlike mammals, the *Drosophila* genome encodes a single member of the NFAT gene family, which does not possess calcineurin binding sites, and is therefore insensitive to intracellular Ca^2+^ and SOCE (Keyser et al., 2007).

In order to identify transcription factors (TFs) that regulate SOCE-mediated gene expression in *Drosophila* neurons, we examined the upstream regions (up to 10 kb) of SOCE-regulated genes using motif enrichment analysis (Zambelli et al., 2009) for TF binding sites (Figure 4A). This analysis helped identify several putative TFs with enriched binding sites in the regulatory regions of SOCE-regulated genes (Figure 4B, Figure 4-figure supplement 1) which were then analyzed for developmental expression trajectories (Hu et al., 2017) in the *Drosophila* CNS (Figure 4B; lower panel). Among the top 10 identified TFs, the homeobox transcription factor Trl/GAF had the highest enrichment value, lowest p-value (Figure 4B) and was consistently expressed in the CNS through all developmental stages, with a distinct peak during pupal development (Figure 4B; lower panel). To test the functional significance of Trl/GAF, we tested flight in flies with *THD’* specific knockdown of *Trl* using two independent RNAi lines (Figure 4C) as well as in existing *Trl* mutant combinations that were viable as adults (Figure 4-figure supplement 2). While homozygous *trl* null alleles are lethal, trans-heterozygotes of two different hypomorphic alleles were viable and exhibit significant flight deficits (Figure 4C, Figure 4-figure supplement 2). Moreover, the flight deficits caused by *Trl^RNAi^* specifically in the *THD’* neurons could be rescued by *Set2* overexpression (Figure 4C; compare data in red and brown).

**Figure 4.**
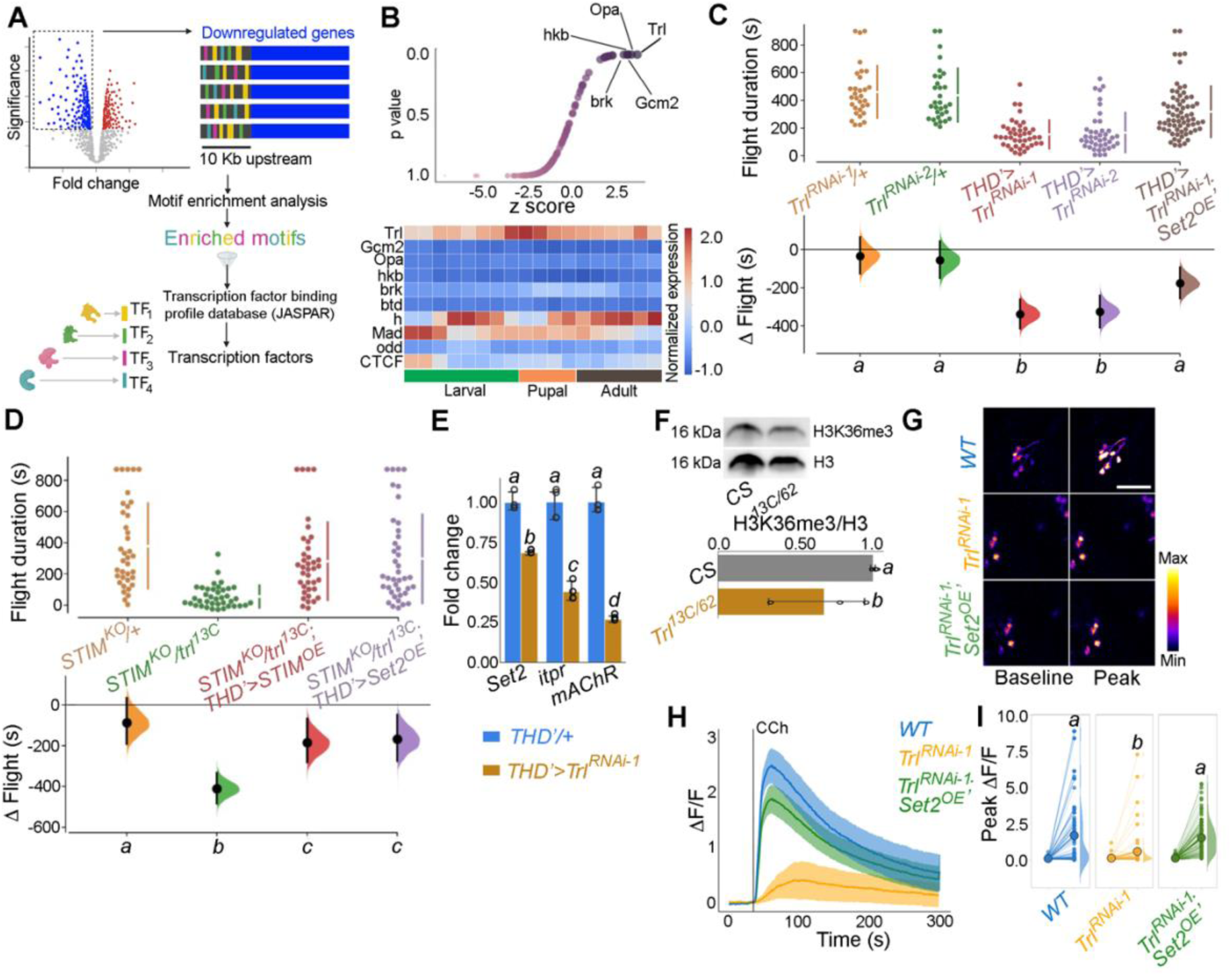
Identification of Trl as an SOCE responsive transcription factor. 1. Schematic of motif enrichment analysis for identification of putative SOCE dependent transcription factors (TFs). **B.** Candidate TFs identified (upper panel) and their expression through development (lower panel; modEncode(modENCODE Consortium et al., 2010)) suggests Trl as a top SOCE-responsive candidate TF. **C**. Genetic depletion of *Trl* in *THD’* DANs results in significant flight defects, that can be rescued by overexpression of *Set2*. **D.** Heteroallelic combination of the *Trl* hypomorphic allele (*trl^13C^*) with a *STIM* deficiency causes significant flight deficits, that can be rescued by overexpression of *STIM* or *Set2*. **E.**qRT-PCRs measured relative to *rp49* show reduced *Set2, itpr, and mAChR* levels in *THD’* DANs upon *Trl* knockdown with *Trl*^RNAi-1^. Individual data points of 4 biological replicates are shown as circles and mean expression level as a bar plot (± SEM). The letters above indicate statistically indistinguishable groups after performing a Kruskal Wallis Test and a Mann-Whitney U-test (p<0.05). **F.** A representative western blot (left) showing reduced H3K36me3 levels in a *Trl* mutant combination. Quantification of H3K36me3 from 3 biological replicates of *WT* (control) and *Trl* mutant brain lysates (panel on the right). Letters indicate statistically indistinguishable groups after performing a two-tailed t-test (p<0.05).**G**. Knockdown of *Trl* in *THD’* neurons attenuate Ca^2+^ response to CCh as shown in representative images of GCamP6m fluorescence quantified in (**H-I**). Scale bar = 10 μM*. Set2* overexpression in the background of *THD’>Trl^RNAi^* rescues the cholinergic response (**G-I**). Quantification of Ca^2+^ responses are from 10 or more brains per genotype and were performed as described in the legend to Figure 3. The letters above indicate statistically indistinguishable groups after performing a Kruskal Wallis Test and a Mann-Whitney U-test (p<0.05).

To further test Trl function, as an effector of intracellular Ca^2+^ signaling and SOCE, we investigated genetic interactions of *trl^13C^*, a hypomorphic mutant recessive *Trl* allele, with an existing deficiency for the ER-Ca^2+^ sensor STIM (*STIM^KO^*), an essential activator of the Orai channel (Wang et al., 2010) as well as existing mutants for an intracellular ER-Ca^2+^ release channel, the IP_3_R (Joshi et al., 2004). While *STIM^KO^/+* flies demonstrate normal flight bout durations, adding a single copy of the *trl^13C^* allele to the *STIM^KO^* heterozygotes (*STIM^KO^/+; trl^13C^/+* trans-heterozygotes) resulted in significant flight deficits, which could be rescued by overexpression of either *STIM* or *Set2* (Figure 4D). The rescue of *STIM^KO^/+; trl^13C^/+* with *STIM*^OE^ (Figure 4D) indicates that SOCE, driven by *STIM^OE^*, activates residual Trl encoded by a copy of *Trl^+^* in this genetic background (*trl^13c^/+)* to rescue flight. The role of Trl as an SOCE-regulated transcription factor is further supported by rescue of flight in *STIM^KO^/+; trl13c/+* flies by overexpression of *Set2* (Figure 4D). Set2 expression in *THD’* neurons was demonstrated earlier as requiring Orai mediated Ca^2+^ entry (Figure 2B-C). These genetic data are consistent with the positive feedback loop proposed earlier (Figure 3F). We hypothesize that Trl is non-functional upon expression of *Orai^E180A^* due to reduced SOCE. Loss of Trl function in turn down-regulates *Set2* expression. A single copy of *trl^13C^* placed in combination with a single copy of various *itpr* mutant alleles, also showed a significant reduction in the duration of flight bouts (Figure 4-figure supplement 3), further supporting a role for intracellular Ca^2+^ signaling in Trl function. Flight deficits observed upon specific expression of *Trl^RNAi^* in *THD’* neurons (Figure 4C) indicate the direct requirement of Trl in fpDANs. Taken together, these findings provide good genetic evidence for Trl as a SOCE-responsive TF in *THD’* neurons. Due to the strong flight deficit observed by expression of *Orai^E180A^* we were unable to test genetic interactions of *Trl* with Orai directly. *Orai^RNAi^* strains exhibit off-target effects and *Orai* hypomorphs are unavailable.

The requirement of *Trl* for SOCE-dependent gene expression was tested directly by measuring *Set2* transcripts in FACS sorted *THD’* neurons with knockdown of *Trl* (*THD’>Trl^RNAi^*). *Set2* has 20 Trl binding sites in a 2 Kb region upstream of the transcription start site (Figure 4-figure supplement 4) and knockdown of *Trl* resulted in downregulation of *Set2* (32%) and the Set2 regulated genes *itpr* (56%) and *mAchR* (73%; Figure 4E). Moreover, a trans-heterozygotic combination of hypomorphic Trl alleles (*trl^13C/62^*) had markedly reduced levels of brain H3K36me3 (Figure 4F).

To test if Trl drives cellular function in the fpDANs, we stimulated the mAChR with CCh and measured Ca^2+^ responses in fpDANs with knockdown of *Trl*. Compared to *WT* fpDANs, which respond robustly to cholinergic stimulation, *Trl^RNAi^* expressing DANs, exhibit strongly attenuated responses (Figure 4G-I). Overexpression of *Set2* in the *Trl^RNAi^* background rescued the cholinergic response (Figure 4G-I). The rescue of both flight (Figure 4C, D) and the cholinergic response (Figure 4G-I) by overexpression of *Set2* in flies with knockdown of *Trl* in *THD’* neurons (*THD’>Trl^RNAi^*), confirms that Trl acts upstream of *Set2* to ensure optimal neuronal function and flight.

The results obtained so far indicate that Trl functions as an intermediary transcription factor between SOCE and its downstream effector gene *Set2*. Overexpression of *Trl*, however, is unable to rescue the loss of flight caused by loss of SOCE (*THD’>Orai^E180A^*; Figure 5A). Moreover, *Trl* transcript levels are unchanged in the *Orai^E180A^* condition (Figure 5B). These data could either mean that phenotypes observed upon loss of Trl are independent of Orai-Ca^2+^ entry or that Trl requires Ca^2+^ influx through SOCE for its function to go from an ‘inactive’ form to an ‘active’ form (schematized in Figure 5C). We favor the latter interpretation because key transcripts downregulated by expression of *Orai^E180A^* (*Set2, itpr* and *mAchR*) were also downregulated upon knockdown of *Trl* in *THD’* neurons (see Figures 3E and 4E).

**Figure 5.**
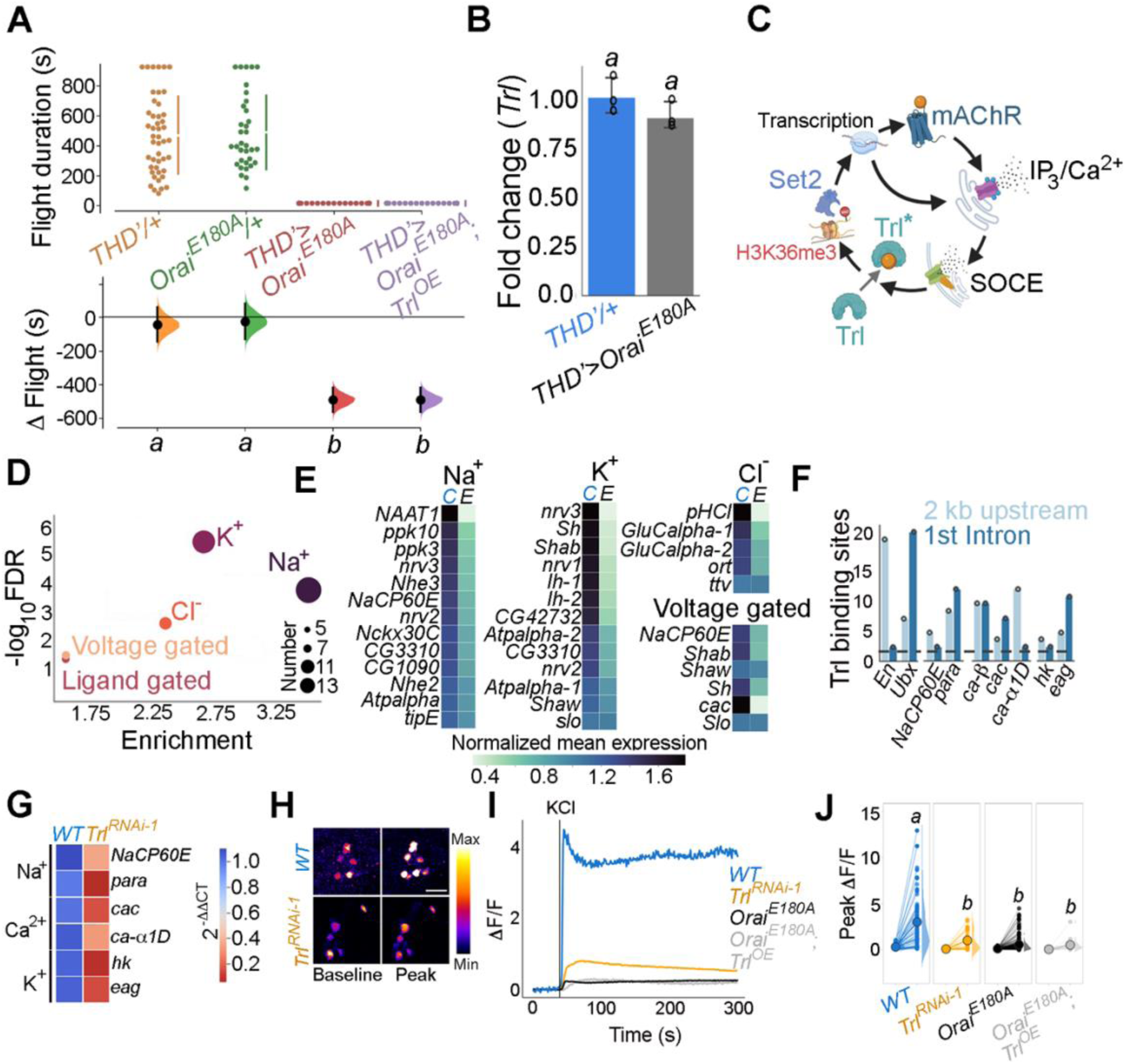
Trl activity downstream of SOCE is required for *THD’* DAN activity. A. Overexpression of *WT Trl* is insufficient to rescue flight deficits in *THD’>Orai^E180A^* flies as evident from flight bout measurements in the indicated genotypes. Flight assay measurements are as described earlier. **B**. *Trl* transcript levels are not altered in *THD’>Orai^E180A^* neurons with loss of SOCE. qRT-PCR data are measured relative to *rp49*. The bar plot indicates mean expression levels, with individual data points represented as circles. The letters above indicate statistically indistinguishable groups from 3 independent biological replicates after performing a two-tailed t-test (p<0.05). **C**. Schematic with possible Ca^2+^ - mediated activation of Trl downstream of Orai-mediated Ca^2+^ entry (SOCE). **D.** Genes encoding ion channels are enriched among SOCE responsive genes in *THD’* DANs as determined by GO analysis. Circles of varying radii are scaled according to the number of genes enriched in that category. **E**. Downregulation of individual ion channel genes depicted as a heatmap. **F**. Number of Trl binding sites (GAGA repeats) shown as a bar plot in the regulatory regions (2 kb upstream or 1^st^ intron) of key SOCE-responsive ion channel genes compared to known Trl targets (*En* and *Ubx*). The dashed line indicates the average expected number of Trl targets. **G**. Heatmap of 2^-ΔΔCT values measured using qRT-PCRs from sorted *THD’* DANs with knockdown of *Trl*. **H.** Representative images of KCl-induced depolarizing responses in *THD’* DANs with knockdown of *Trl*. (Scale bar = 10 μM) quantified in **I and J**. Quantification of Ca^2+^ responses are from 10 or more brains per genotype and were performed as described in the legend to Figure 3. The letters above indicate statistically indistinguishable groups after performing a Kruskal Wallis Test and a Mann-Whitney U-test (p<0.05).

To understand how Orai-mediated Ca^2+^ entry might activate Trl we went back to previously known biochemical characterization of Trl. Although Trl does not possess a defined Ca^2+^ binding domain, it interacts with a diverse set of proteins, primarily through its BTB-POZ domain (Figure 5-figure supplement 1) as demonstrated earlier by affinity purification of Trl from embryonic extracts followed by high throughput mass spectrometry (Lomaev et al., 2017). Among the identified interacting partners, we focused on kinases, keeping in mind an earlier study in *Drosophila* indicating phosphorylation of Trl at a threonine residue (T237; Zhai et al., 2008). Analysis of the Trl interactome revealed several kinases, which were expressed to varying extents in the *THD’* neurons (Figure 5-figure supplement 2), including the Ca^2+^ dependent CamKII, implicated earlier in flight in *THD’* neurons (Ravi and Hasan 2018). Loss of CamKII in *THD’* neurons by RNAi mediated knockdown or through expression of a peptide inhibitor (Ala; Mehren and Griffith, 2004) resulted in significant flight deficits (Figure 5-figure supplement 3). To test if CamKII activation downstream of SOCE is required for flight, we expressed a constitutively active version of CamKII (*T287D*; Kadas et al., 2012) in the background of *THD’*>*Orai^E180A^*. The phosphomimetic *CamKII^T287D^* point mutation renders CamKII activity Ca^2+^ independent (Malik and Hodge, 2014). Acute expression of *CamKII^T287D^* in the *Orai^E180A^* background using the TARGET system (McGuire et al., 2004) resulted in a weak rescue of flight when implemented from 72 to 96 hrs APF (Figure 5-figure supplement 4) whereas overexpression of WT *CamKII* or a phosphorylation-incompetent allele (*CamKII^T287A^*), failed to rescue flight (Figure 5-figure supplement 4). These data support a model (schematized in Figure 5C) wherein Ca^2+^ influx through SOCE sustains Trl activation, in part through CamKII, leading to expression of *Set2* followed by increased levels of chromatin H3K36me3 marks that drive further downstream gene expression changes through a transcriptional feedback loop. Our data do not rule out activation of other Ca^2+^ sensitive mechanisms for activation of Trl. Moreover, rescue of flight in Trl mutant/knockdown conditions by *STIM* overexpression (Figure 4C and 4D) suggests the presence of additional SOCE-responsive transcription factors in fpDANs.

### Trl function downstream of SOCE targets neuronal activity through changes in ion channel gene expression including VGCCs

Next we investigated the extent to which the identified SOCE-Trl-Set2 mechanism impacts fpDAN function. ‘Ion transport’ is among the top GO terms identified in SOCE-responsive genes (Figure 2A). Indeed, multiple classes of ion channel genes including several Na^+,^ K^+^ and Cl^-^ channels are downregulated in fpDANs upon expression of *Orai^E180A^* (Figure 5D, E). Regulation of ion channel gene expression by Trl was indicated from the enrichment of Trl binding sites in regulatory regions of some ion channel loci (Figure 5F). We purified fpDANs with knock down of *Trl* (*THD’>Trl^RNAi-1^*) and measured expression of a few key voltage-gated ion channel genes (Figure 5F). *NaCP60E*, *para* (Na^+^ channels), *cac*, *ca-α1D* (VGCC subunits), *hk*, and *eag* (outward rectifying K^+^ channels) are all downregulated between 0.6-0.9 fold upon *Trl* knockdown (Figure 5G) as well as upon loss of SOCE (Figure 5B). Taken together, these results indicate that the expression of key voltage-gated ion channel genes which are required for the depolarization-mediated response, maintenance of electrical excitability and neurotransmitter release is a focus of the transcriptional program set in place by SOCE-Trl-Set2.

To test the functional consequences of altered ion channel gene expression downstream of Trl, neuronal activity of the fpDANs was tested by means of KCl-evoked depolarisation. PPL1 DANs exhibited a robust Ca^2+^ response that was lost upon knock down of *Trl* (Figure 5H-J; data in orange). In consonance with the earlier behavioural experiments (Figure 5A), overexpression of *Trl^+^* in *THD’* DANs did not rescue the KCl response in PPL1 DANs with loss of SOCE (*OraiE^180A^; Trl^OE^*, Figure 5I, J), further reinforcing the hypothesis that, Trl activity in fpDANs is dependent of Ca^2+^ entry through Orai.

Response to KCl is also lost in PPL1 neurons by knockdown of *Set2* and restored to a significant extent by overexpression of *Set2* in *THD’* DANs lacking SOCE (*Orai^E180A^;Set2^OE^*^;^ Figure 6A-C). We confirmed that the KCl response of PPL1 neurons requires VGCC function, either by treatment with Nimodopine (Xu and Lipscombe, 2001), an L-type VGCC inhibitor or by knockdown of a conserved VGCC subunit *cac (cac^RNAi^)*. Both forms of perturbation abrogated the depolarisation response to KCl in PPL1 neurons (Figures 6D-F; data in purple or orange respectively). Moreover, the Ca^2+^ entry upon KCl-mediated depolarization in PPL1 neurons is a cell-autonomous property as evident by measuring the response after treatment with the Na^+^ channel inhibitor, Tetrodotoxin (TTX; Figures 6D-F; data in magenta).

**Figure 6.**
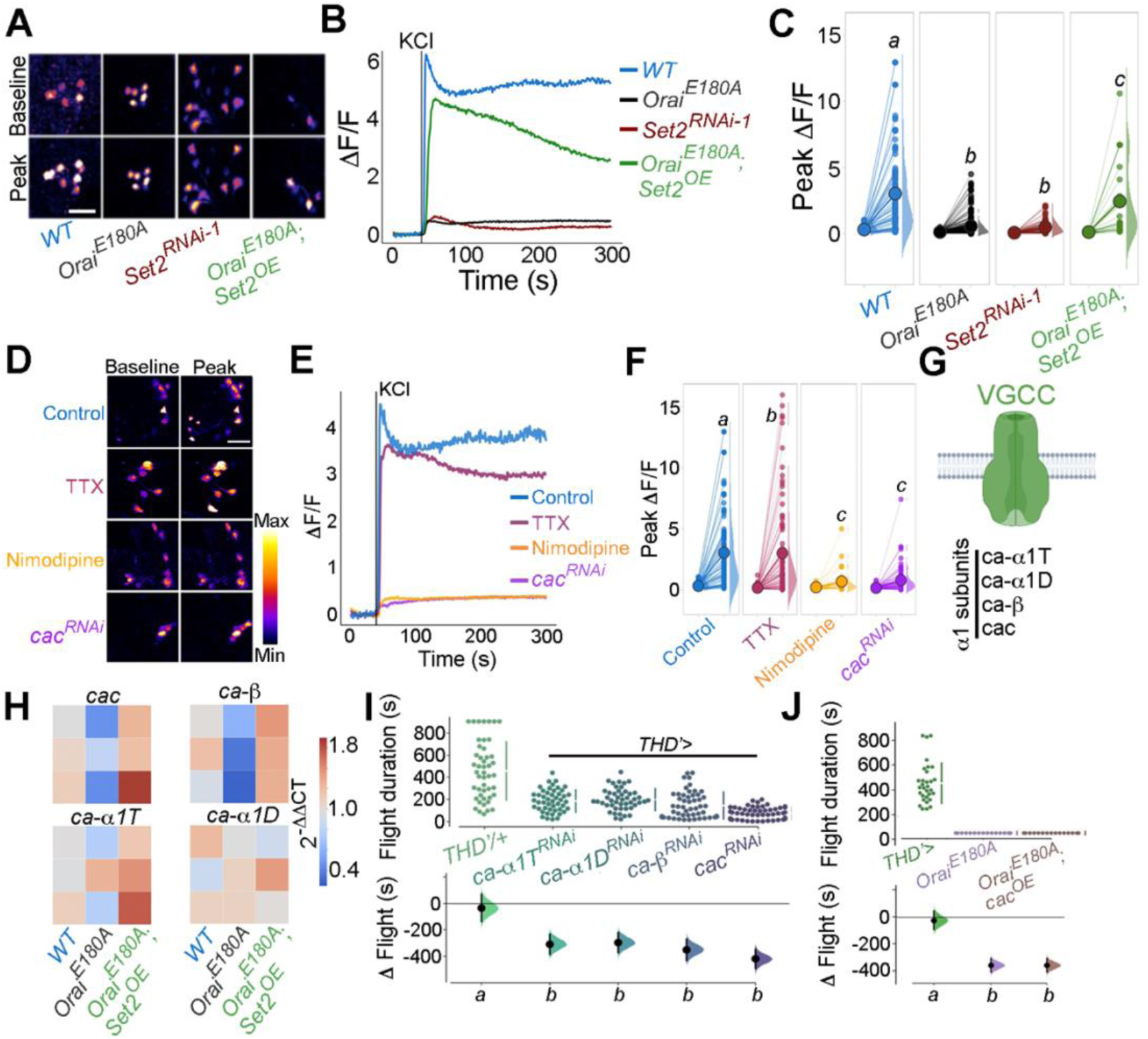
fpDAN excitability requires Orai-mediated Ca^2+^ entry acting through *Set2* -mediated VGCC gene expression. KCl-induced depolarizing responses in *fp*DANs of the indicated genotypes measured using GCaMP6m indicate a requirement for Orai and *Set2*. Representative images **A. (**scale bar = 10 μM), quantified in **B.** and **C**. KCl evoked responses are modulated upon treatment with 10 μM TTX (magenta), 10 μM Nimodipine (orange), and upon *cac^RNA^*^i^ (purple)-representative images (scale bar = 10 μM), (**D**), quantified in **E**. Median KCl-evoked GCaMP6m responses plotted as a function of time. The solid line indicates the time point of addition of 70 mM KCl. KCl responses are quantified as a paired plot of peak responses and **F**. with letters representing statistically indistinguishable groups as measured using a Kruskall Wallis test and post hoc Mann-Whitney U test (p<0.05). Quantification of Ca^2+^ responses are from 10 or more brains per genotype and were performed as described in the legend to Figure 3. **G.** Schematic of a typical VGCC and its constituent subunits. **H.** Heatmap of 2^-ΔΔCT values measured using qRT-PCRs from sorted *THD’* DANs show a reduction in the expression of VGCC subunit genes upon loss of Orai function and a rescue by *Set2* overexpression. **I**. Flight assays representing the effect of various VGCC subunit gene RNAis showing flight defects to varying extents. Overexpression of the key VGCC subunit gene-*cac* is required (**I**) but not sufficient (**J**) for restoring flight defects caused by loss of *Orai* function. Flight assay measurements are as described earlier.

Having confirmed that the Ca^2+^ response towards KCl depolarisation requires VGCCs, we tested the expression level of key components of *Drosophila* VGCCs (Figure 6G) in *THD’* DANs, including *cacophony* (*cac*), *ca-α1D*, *ca-α1T*, and *ca-β*. All four subunits of *Drosophila* VGCC are downregulated upon loss of SOCE (*Orai^E180A^*; Figure 6H) in *THD*’ DANs whereas overexpression of *Set2* in the background of *Orai^E180A^* restored expression of the four VGCC subunits tested (Figure 6H), thus explaining recovery of the KCl response upon *Set2* overexpression in PPL1 neurons lacking SOCE (*Orai^E180A^*; Figures 6A-C).

The functional significance of downregulation of VGCC subunits by the SOCE-Trl-Set2 pathway was tested directly by knockdown of the 4 VGCC subunits independently in PPL1 neurons followed by measurement of flight bout durations. Significant loss of flight was observed with knockdown of each subunit (Figure 6I) though in no case was the phenotype as strong as what is observed with loss of SOCE (*Orai^E180A^*; Figure 1D). These data suggest that in addition to VGCC subunit expression, appropriate expression of other ion channels in the fpDANs is of functional significance. This idea is further supported by the observation that loss of flight upon loss of SOCE cannot be restored by overexpression of *cac* alone (Figure 6J) indicating that optimal neuronal activity requires an ensemble of genes, including ion channels, whose expression is regulated by SOCE-Trl-Set2.

### *THD’* DANs require SOCE for developmental maturation of neuronal activity

The functional relevance of changes in neuronal activity during pupal maturation of the PPL1-dependent flight circuit was tested next. Inhibition of *THD’* DANs from 72 to 96 hours APF using acute induction of the inward rectifying K^+^ channel *Kir2.1*(Johns et al., 1999) or optogenetic inhibition through the hyperpolarizing Cl^-^ channel *GtACR2* (Govorunova et al., 2015) resulted in significant flight deficits (Figure 7A; data in dark or light green). Similar flight defects were recapitulated upon hyperactivation of *THD’* neurons through either optogenetic (*CsChrimson*; Klapoetke et al., 2014) or thermogenetic (*TrpA1*; Viswanath et al., 2003) stimulation (Figure 7A; data in orange or red), indicating that these neurons require balanced neuronal activity during a critical window in late pupal development, failing which the flight circuit malfunctions. To test if restoring excitability in fpDANs lacking SOCE *(THD’>Orai^E180A^*) is sufficient to restore flight, we induced hyperexcitability by overexpressing *NachBac* (Nitabach, 2006) a bacterial Na^+^ channel. *NachBac* expression rescued both flight (*THD’>Orai^E180A^*; Figure 7B) and excitability (Figures 7C-E). A partial rescue of flight was also obtained by optogenetic stimulation of neuronal activity through activation of *THD’>Orai^E180A^* DANs using *CsChrimson* either 72-79 hrs APF or 0-2 days post eclosion (Figure 7E). In this case, flight rescues were accompanied by a corresponding rescue of *CsChrimson* activation induced Ca^2+^ entry (Figures 7F, G). Moreover inducing neuronal hyperactivation with indirect methods such as excess K^+^ supplementation (Figure 7-figure supplement 1) or impeding glial K^+^ uptake, by genetic depletion of the glial K^+^ channel *sandman* (Weiss et al., 2019; Figure 7-figure supplement 2), partially rescued flight in animals lacking SOCE in the *THD’* DANs. Together, these results indicate that SOCE, through Trl and Set2 activity, determines activity in fpDAN during circuit maturation by regulating expression of voltage-gated ion channel genes, like *cac* (see discussion).

**Figure 7.**
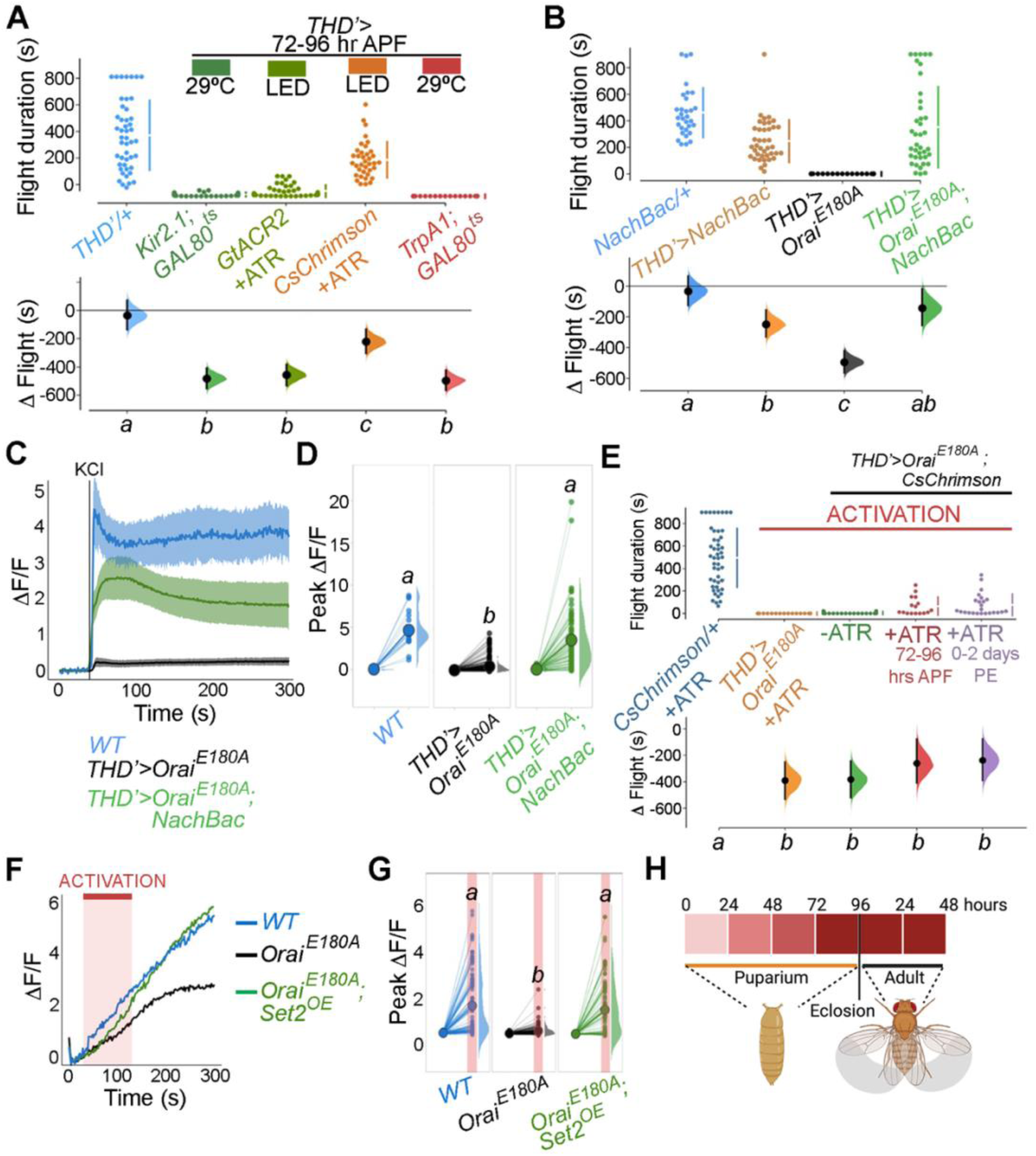
SOCE-mediated gene expression sets the excitability threshold during pupal development. **A.** Altering excitability in *THD’* DANs using *Kir2.1* or *GtACR2* mediated inhibition or *CsChrimson* or *TrpA1* during the critical 72-96 hour APF developmental window results in significant flight defects. Orai loss of function phenotypes can be rescued by overexpression of *NachBac* or *CsChrimson* in terms of flight bout durations (**B, E**) and depolarizing KCl responses (**C, D, D, G**). **H**. Schematic for Orai-mediated Ca^2+^ entry regulating expression of key genes that regulate the excitability threshold of dopaminergic neurons regulating flight during a critical developmental window. Flight assays are represented as described earlier, (n>30). Ca^2+^ responses were quantified as described in the legend to Figure 3 and were from 10 or more brains per genotype. Letters above each genotype represent statistically indistinguishable groups as measured using a Kruskall-Wallis test and post hoc Tukey test (p<0.05).

fpDANs also require a balance between H3K36me3/H3K27me3 mediated epigenetic regulation (Figure 2K). To understand if this epigenetic balance affects flight through modulation of neuronal excitability, we measured KCl-induced depolarizing responses of PPL1 DANs in Orai deficient animals (*THD’>Orai^E180A^*) fed on the H3K27me3 antagonist GSK343 (0.5 mM). GSK343 fed animals were sorted into 2 groups (Fliers/Non-Fliers) on the basis of flight bout durations of greater or lesser than 30 s (Figure 7-figure supplement 3A), and tested for responses to KCl (Figure 7-figure supplement 3B, C). *THD’>Orai^E180A^* flies fed on GSK343 showed a rescue in KCl responses, with a clearly enhanced response in the fliers compared to the non-fliers (Figure 7-figure supplement 3B, C), thereby reiterating the hypothesis that SOCE-mediated regulation of activity in these neurons acts through epigenetic regulation between opposing histone modifications.

### *THD’* DANs require SOCE-mediated gene expression for axonal arborization and neuromodulatory dopamine release in the mushroom body γ lobe

Next, we re-visited how loss of activity during the critical maturation window of 72-96 hrs APF and 0-2 days post-eclosion affects axonal arborisation of fpDANs that innervate the γ lobe. From previous work we know that major axonal branches of the fpDANs reach the γ lobe normally in *THD’>Orai^E180A^* animals (Figure 8A, B; Pathak et al., 2015). Neuronal activity can drive changes in neurite complexity and axonal arborization (Depetris-Chauvin et al., 2011) especially during critical developmental periods (Sachse et al., 2007). To understand if Orai mediated Ca^2+^ entry and downstream gene expression through Set2 affects this activity-driven parameter, we investigated the complexity of fpDAN presynaptic terminals within the γ2α’1 lobe MB using super-resolution microscopy (Figure 8 A, B). Striking changes in the neurite volume upon expression of *Orai^E180A^* were observed. Importantly these changes could be rescued to a significant extent by restoring either *Set2* (*Orai^E180A^; Set2^OE^*) or by inducing hyperactivity through *NachBac* expression (*Orai^E180A^; NachBac^OE^*; Figure 8 C, D).

**Figure 8.**
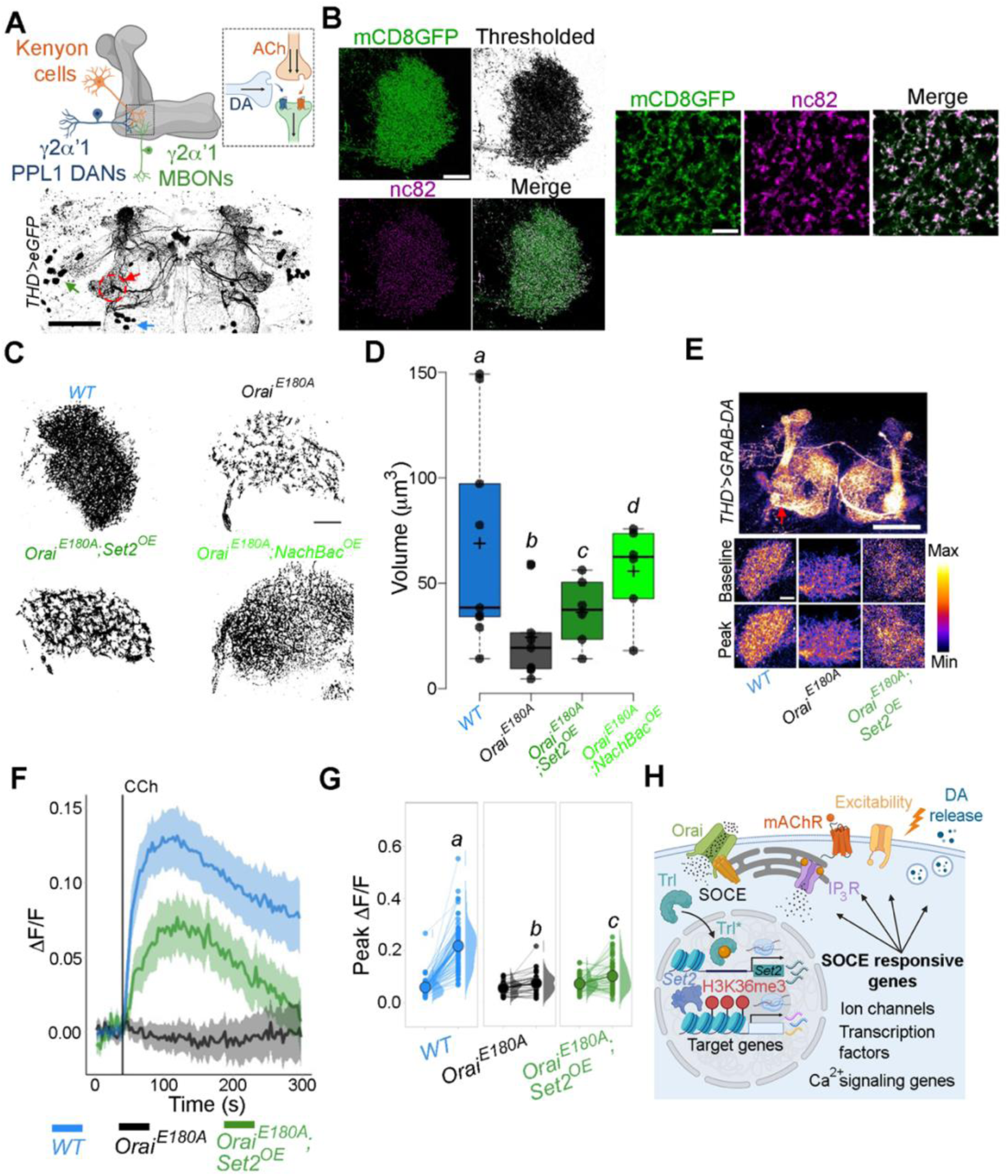
*THD’* DANs require SOCE-mediated gene expression for axonal arborization and neuromodulatory dopamine release in the mushroom body γ lobe. **A.** Schematic of a KC-DAN-MBON tripartite synapse at the γ2α’1 MB lobe (upper), innervated by fpDANs (lower). The γ2α’1 MB lobe is marked by a red circle and arrow and the fpDAN clusters are marked with a green arrow (PPL1) and blue arrow (PPM3). Scale bar=50 μM. **B**. Representative images of the mushroom body γ lobe marked with THD’ driven mCD8GFP (green) and immunolabelled with anti-nc82 (Brp) antibody (magenta) to mark the presynaptic terminals. Scale bar = 5 μm. **C**. Representative optical sections through the mushroom body γ lobe from the indicated genotypes. Changes in axonal arborization observed were quantified as total projection volume in **D**. **E**. CCh-evoked DA release measured at the γ2α’1 MB lobe using the GRAB-DA sensor expressed in *THD’* DANs. Representative GRAB-DA images of *THD’* DANs are shown with baseline and peak evoked responses in the indicated genotypes. Scale bar = 10 μM. **F**. Median GRAB-DA responses plotted as a function of time. A shaded region around the solid line represents the 95% confidence interval from 10 or more brains per genotype. **G.** Individual cellular responses depicted as a paired plot of peak responses where different letters above indicate statistically distinguishable groups after performing a Kruskal Wallis Test and a Mann-Whitney U-test. **H**. Schematic describing how Orai-mediated Ca^2+^ entry regulates expression of key genes that control neuronal activity in fpDANs.

To understand if reduced axonal arborisation within the γ lobe has functional consequences we measured CCh-evoked DA release at the γ2α’1 region of the mushroom body (Figure 8A; red arrow). For this purpose we used the dopamine sensor *GRAB-DA* (Sun et al., 2018). Loss of SOCE (*THD’>Orai^E180A^*) attenuated DA release (Figures 8E, F, G), which could be partially rescued by overexpression of *Set2*. Taken together these data identify an SOCE-dependent gene regulation mechanism acting through the transcription factor Trl, and the histone modifier Set2 for timely expression of genes that impact diverse aspects of neuronal function including neuronal activity, axonal arborisation, and sustained neurotransmitter release (schematised in Figure 8H).

## Discussion

Over the course of development, neurons define an excitability set point within a dynamic range (Truszkowski and Aizenman, 2015), which stabilizes existing connections (Mayseless et al., 2023) and enables circuit maturation (Johnson-Venkatesh et al., 2015). The setting of this threshold for individual classes of neurons is based on the relative expression of various ion channels. In this study, we identify an essential role for the store-operated Ca^2+^ channel *Orai* in determining ion channel expression and neuronal activity in a subset of dopaminergic neurons central to a MB circuit for *Drosophila* flight. Orai*-*mediated Ca^2+^ entry, initiated by neuromodulatory acetylcholine signals, is required for amplification of a signalling cascade, that begins by activation of the homeo-box transcription factor Trl, followed by upregulation of several genes, including *Set2*, a gene that encodes an enzyme for an activating epigenetic mark, H3K36me3. Set2 in turn establishes a transcriptional feedback loop to drive expression of key neuronal signaling genes, including a repertoire of voltage-gated ion channel genes required for neuronal activity in mature PPL1 dopaminergic neurons (schematized in Figure 9).

**Figure 9.**
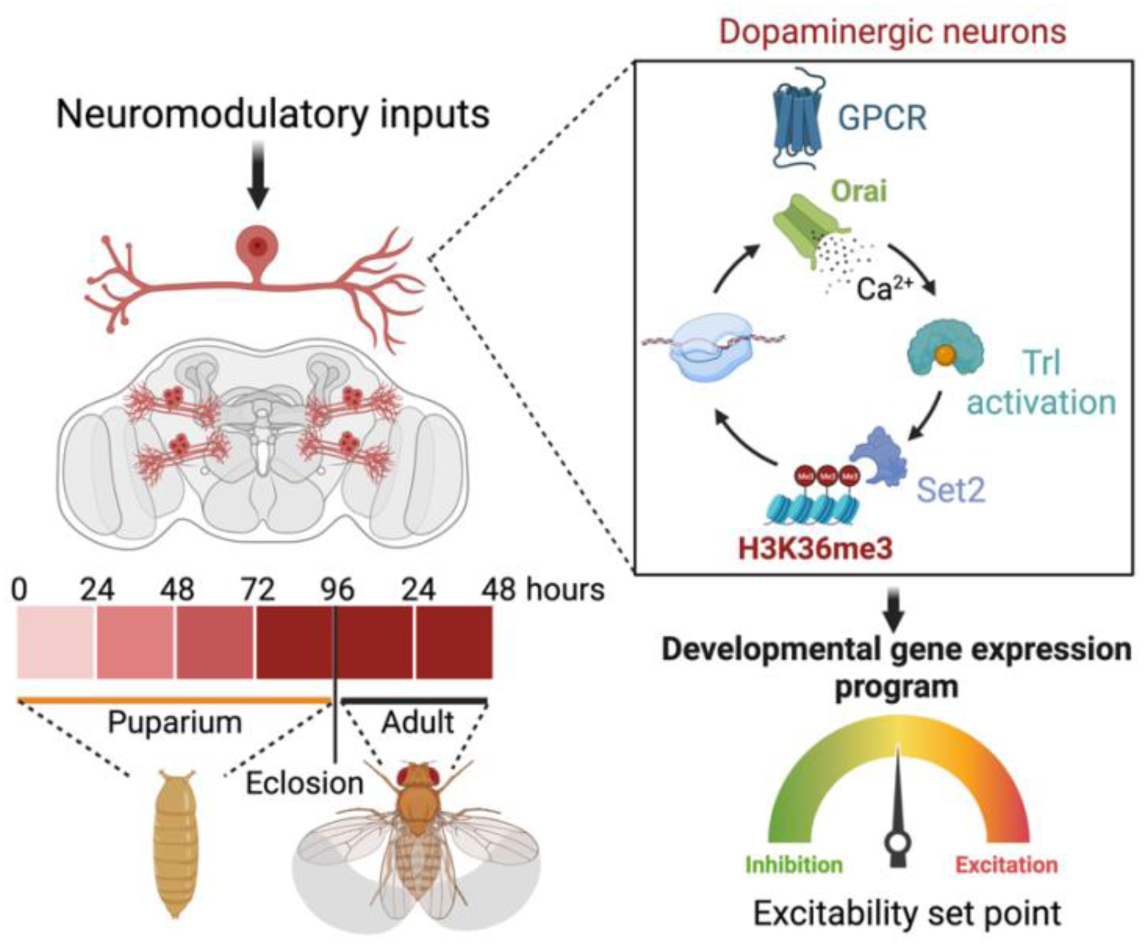
Schematic outline of the role of SOCE in determining excitability and function of dopaminergic neurons required for flight circuit development. SOCE acts through a transcriptional feedback loop including the transcription factor Trl, and the H3K36 methyltransferase Set2 to induce the expression of genes required for neuronal excitability and flight.

### SOCE supports gene expression transitions during critical developmental windows

Studies in the *Drosophila* mushroom body have identified spontaneous bouts of voltage-gated Ca^2+^ channel mediated neuronal activity, through as yet unknown mechanisms, which occur during early adulthood driving refinement and maturation of behaviours such as associative learning (Leinwand and Scott, 2021). Our studies here, on a subset of 21-23 central dopaminergic neurons (Figures 1B, C) of which some send projections to the mushroom body and regulate flight behaviour, provide a mechanistic explanation for neuromodulation-dependent intracellular signaling culminating in transcriptional maturation of a circuit supporting adult flight. In the critical developmental window of 72-96 hrs APF, a cohort of SOCE-responsive genes, that include a range of ion channels, undergo a wave of induction. Loss of SOCE at this stage thus renders these neurons functionally incompetent (Figures 3B-D, 6G-I, 7D-F) and abrogates flight (Figure 1D). Developmentally assembled neuronal circuits require experience/activity-dependent maturation (Akin and Zipursky, 2020). The requirement of SOCE for flight circuit function during the early post eclosion phase (0-2 days) suggests that maturation of the flight circuit extends into early adulthood, where presumably feedback from circuit activity may further refine synaptic strengths (Sugie et al., 2018). Expression of *Orai^E180A^* in 5 day old adults had no effect on flight (Figure 1E), indicating that either SOCE is not required for maintaining gene expression in adults or that wildtype Orai channel proteins from late pupal and early adults perdure for long periods extending into adulthood. Loss of SOCE in adult brains beyond 5-7 days, when Orai protein turnover might be expected, has not been assessed so far. The absence of any visible changes to primary neurite patterning suggests that SOCE in the MB-DANs works primarily to refine synaptic function at pre-synaptic terminals within the MB lobe. Taken together, our findings, identify a role for SOCE in restructuring the neuronal transcriptome during a critical developmental transition, which facilitates key changes in neuronal function required for circuit formation and adult behaviour (schematised in Figure 9). In summary, our findings here identify Orai-driven Ca^2+^ entry as a key signaling step for driving coordinated gene expression enabling the maturation of nascent neuronal circuits and sustenance of adult behaviour.

### SOCE regulates a balance between competing epigenetic signatures

Chromatin structure and function is dynamic over the course of brain development (Kishi and Gotoh, 2018). One way neurons achieve dynamic spatio-temporal control over developmental gene expression is via epigenetic mechanisms such as post translational histone modifications (Geng et al., 2021). Cells possess an extensive toolkit of histone modifiers, with characterised effects on transcriptional output by either activating or repressing gene expression (Bannister and Kouzarides, 2011). However, relatively little, is known about how modifiers of histone marks are in turn regulated to bring about the requisite changes in neuronal gene expression over developmental timescales.

The SET-domain containing family of histone modifiers (Figure 2B), including H3K36me3 methyltransferase Set2 (Figure 2C), identified here appear to function as key transcriptional effectors induced downstream of neuromodulatory inputs that stimulate Store-operated Ca^2+^ entry through Orai during pupal and adult maturation of flight promoting dopaminergic neurons. SOCE-driven *Set2*-mediated H3K36me3 enhances the expression of key GPCRS, components of intracellular Ca^2+^ signaling (Figure 2A) and ion channels (Figures 6A, B) for optimal dopaminergic neuron function (Figure 7D-F) and flight (Figure 1C). Rescue of flight by genetic depletion of an epigenetic ‘eraser’ *Kdm4B*, an H3K36me3 demethylase (Figure 2D) suggests that a balance between perdurance of H3K36me3 and its removal is actively maintained on the expressed genes. Interestingly, another member of the SET domain family, *E(z)* is up-regulated upon loss of SOCE (Figure 2B)*. E(z)* is a component of the *Drosophila* PRC2 complex (Polycomb repressive complex), which represses gene expression through H2K27me3 (Margueron and Reinberg, 2011). Structural studies on *Drosophila* nucleosomes have elucidated that the H3K36me3 modification allosterically inhibits the PRC2 complex activity (Finogenova et al., 2020). Our findings support a model wherein at key developmental stages, SOCE determines the level of these competing epigenetic signatures thus allowing gene expression by enhancing Set2 mediated H3K36me3 (Figure 2H). Loss of SOCE leads to deficient Set2-mediated H3K36me3 (Figure 2E), a likely shifting of the balance in favour of PRC2 mediated H3K27me3 and overall repression in gene expression (as observed in Figure 1-figure supplements 8-9). Feeding SOCE-deficient animals a pharmacological inhibitor of PRC2 (GSK434) led to a significant rescue in behaviour (Figure 2J) presumably by disinhibition of H3K27me3 mediated gene repression. Though flight initiation is rescued in a dose dependent manner, the duration of flight bouts is not sustained (Figure 2-figure supplement 8) indicating that suppression of H3K27me3 marks in the SOCE-deficient flies is partial. These findings indicate additional roles for other histone modifications such as Set1 mediated H3K4me3, which may work in concert to regulate developmental gene expression programs stimulated by SOCE. Future studies directed at a comprehensive cataloguing of multiple epigenetic signatures in defined neuronal subsets using cell-specific techniques should help elucidate these mechanisms.

### Trl as an SOCE responsive transcription factor

SOCE-mediated regulation of gene expression has been reported in mammalian neural progenitor cells (Somasundaram et al., 2014) and *Drosophila* pupal neurons (Pathak et al., 2015; Richhariya et al., 2017), through as yet unknown mechanisms. A combination of motif enrichment analysis over regulatory regions of SOCE-responsive genes and expression enrichment analysis in cells of interest, helped generate a list of putative SOCE-responsive transcription factors (Figure 4A, B). Among these we experimentally validated Trl/Trithorax-like/GAGA factor (GAF) as a transcription factor required for maturation of excitability in flight promoting dopaminergic neurons using genetic tools (Figure 4). We found that loss of *Trl* in the fpDANs results in significant flight deficits, which could be partially rescued through excess STIM (which raises SOCE). This indicates that increased Ca^2+^ entry through Orai either activates residual Trl or alternate SOCE-responsive transcription factors to induce *Set2* expression and rescue downstream gene expression essential for flight.

Although Trl function has been reported in the context of early embryonic development, where it has been implicated in zygotic genome activation (Gaskill et al., 2021) (ZGA), expression of *Hox* genes (Shimojima, 2003), dosage compensation (Greenberg et al., 2004), and expression of a *Drosophila* voltage-gated calcium channel subunit in germ cell development (Dorogova et al., 2014), this is the first ever report of its role in regulating neuronal gene expression in the context of SOCE and circuit maturation. The molecular mechanism by which Orai-mediated Ca^2+^ entry leads to activation of Trl needs further elucidation. Our finding that CaMKII hyperactivation partially rescues flight deficits caused by loss of Orai function, taken together with the bioinformatic prediction for CaMKII-mediated Trl phosphorylation need to be experimentally verified. Future studies could be directed towards looking at interactions between these two proteins using in vitro biochemical assays.

In vitro studies on *Drosophila* Trl reveal that it utilizes a Q-rich intrinsically disordered domain (IDD) to self-multimerize or interact with a wide range of accessory proteins (Wilkins and Lis, 1999). Multimeric complexes of Trl reside on chromatin with an exceptionally long residence time and maintain chromatin in an ‘open’ state (Tang et al., 2022). Trl has also been reported to directly interact with components of the Transcription initiation and elongation complex (Chopra et al., 2008; Li et al., 2013), support long distance promoter-enhancer interactions (Mahmoudi, 2003), mediate active ATP-dependent chromatin remodeling by maintaining nucleosome free stretches (Tsukiyama et al., 1994), and regulating global gene expression by controlling transcriptional stalling and pausing (Tsai et al., 2016). A recent study demonstrates a role for Trl in chromatin folding in *Drosophila* neurons to enable cell type specific gene expression (Mohana et al., 2023). SOCE-dependent gene regulation may rely upon multiple transcription factors acting in concert in a cell type and developmental stage specific contexts, of which Trl may be just one. Future studies directed at other possible transcription factors downstream of SOCE would be of interest.

### SOCE and the control of neuronal activity and cholinergic neuromodulation of a tripartite synapse in the mushroom body

Neurons undergo maturation of their electrical properties with a gradual increase in depolarizing responses and synaptic transmission over the course of pupal development (Jarvilehto and Finell, 1983). Here we show that a subset of dopaminergic neurons require a balance of ion channels for optimal excitation and inhibition essential for adult function. Neuromodulatory signals, such as acetylcholine are essential for acquiring this balance and act via an SOCE-Trl-Set2 mechanism during late pupal and early adult stages. The ability of *Set2* overexpression to rescue SOCE-deficient phenotypes (Figures 6A-C) indicates that anomalous gene expression is the underlying basis of this phenotype. Indeed, SOCE deficient dopaminergic neurons show reduced expression of an ensemble of voltage-gated ion channel genes (Figures 5E, F) including genes encoding the outward rectifying K^+^ channel *shaw*, slow-inactivating voltage gated K^+^ channel *shaker*, a Ca^2+^ gated K^+^ channel *slowpoke* and *cacophony* a subunit of the voltage gated Ca^2+^ channel (Figure 6G). Ion channels like *Shaker* and *slowpoke* play an important role in establishing resting membrane potential (Singh and Wu, 1990), neuronal repolarisation (Lichtinghagen et al., 1990) and mediating afterhyperpolarisation (AHP) after repeated bouts of activity (Ping et al., 2011). Though dopaminergic neurons require *Set2* mediated expression of *cacophony* for excitability, *cacophony* is in itself insufficient to rescue the loss in excitability and flight deficits upon loss of SOCE (Figure 6J). Future experiments that directly visualise electrophysiological responses of fpDANs from different genotypes would help resolve the role of Cac and other ion channels and determine their individual contributions to THD’ neuronal activity and excitability. Both flight and KCl-depolarisation in SOCE-deficient neurons could, however, be rescued by expression of a heterologous depolarisation activated Na^+^ channel NachBac (Nitabach, 2006) (Figures 7B-D), which increases neuronal excitability by stimulating a low threshold positive feedback loop and to a lesser extent by CsChrimson-mediated optogenetic induction of neuronal activity (Figure 7E). The poorer rescue by optogenetic activation may stem from the fact that sustained bursts of activity are non-physiological and can deplete the readily releasable pool of neurotransmitters (Arrigoni and Saper, 2014; Kaeser and Regehr, 2017). Other indirect forms of inducing neuronal hyperactivation (Figure 7-figure supplements 1-2) in SOCE-deficient neurons also achieved minor rescues in flight. These results identify SOCE as a key driver downstream of developmentally salient neuromodulatory signals for expression of an ion channels suite that enables generation of intrinsic electrical properties and functional maturation of the PPL1 DANs, and the flight circuit.

### Limitations of this study

Direct measurements of SOCE are generally performed in cultured cells by depletion of ER-store Ca^2+^ in ‘0’ Ca^2+^ medium, followed by Ca^2+^ -add back to measure SOCE. In this study we were unable to perform similar SOCE measurements from the fpDANs because they consist of 16-19 neurons in each hemisphere (PPL1 are 10-12 and PPM3 are 6-7 cells; Pathak et al., 2015) and identifying these few neurons was not technically feasible in culture. Measuring SOCE from these neurons *in vivo* was not possible due to the presence of abundant extracellular Ca^2+^ in the brain. Due to these reasons, we have relied upon using Carbachol to elicit IP_3_-mediated Ca^2+^ release and SOCE as a proxy for *in vivo* SOCE. In previous studies we have shown that Carbachol treatment of cultured *Drosophila* neurons elicits IP_3_-mediated Ca^2+^ release and SOCE (Agrawal et al., 2010). Moreover, expression of *Orai^E180A^* completely blocks SOCE as measured in primary cultures of dopaminergic neurons (Pathak et al., 2015) and CCh-induced IP_3_-mediated Ca^2+^ release is tightly coupled to SOCE in *Drosophila* neurons (Venkiteswaran and Hasan, 2009; Chakraborty et al., 2016; Chakraborty et al., 2017). We posit that our measurements of CCh-evoked changes in cellular Ca^2+^ reflect a composite of IP_3_-mediated Ca^2+^ - release and SOCE.

Although we have provided compelling genetic evidence for Trl as an SOCE-responsive transcription factor, the detailed biochemical basis of Trl activation was beyond the scope of this study. We propose phosphorylation by CamKII as a possible mechanism that needs further investigation.

## Methods

### Fly maintenance

*Drosophila* strains were grown on standard cornmeal medium consisting of 80 g corn flour, 20 g glucose, 40 g sugar, 15 g yeast extract, 4 ml propionic acid, 5 ml p-hydroxybenzoic acid methyl ester in ethanol and 5 ml ortho-butyric acid in a total volume of 1 l (ND) at 25°C under a light-dark cycle of 12 h and 12 h. Canton S was used as wild type throughout. Mixed sex populations were used for all experiments. Several fly stocks used in this study were sourced using FlyBase (https://flybase.org), which is supported by a grant from the National Human Genome Research Institute at the US National Institutes of Health (#U41 HG000739), the British Medical Research Council (#MR/N030117/1) and FlyBase users from across the world. The stocks were obtained from the Bloomington Drosophila Stock Centre (BDSC) supported by NIH P40OD018537. All fly stocks used and their sources are listed in Supplementary Table 1a.

### Flight Assays

Flight assays were performed as previously described (Manjila and Hasan, 2018). Briefly, flies aged 3–5 days of either sex were tested in batches of 8–10 flies, and a minimum of 30 flies were tested for each genotype. Adult flies were anaesthetized on ice for 2–3 min and then tethered between at the head-thorax junction using a thin metal wire and nail polish. Post recovery at room temperature for 2-3 min, an air puff was provided as stimulus to initiate flight. Flight duration was recorded for each fly for 15 min. For all control genotypes, GAL4 or UAS strains were crossed to the Wild Type strain, *Canton S*. Flight assays are represented in the form of a swarm plot, wherein each dot represents a flight bout duration for a single fly. The colours indicate different genotypes. The Δ Flight parameter refers to the mean difference for comparisons against the shared CS control which is shown as a Cumming estimation plot(Ho et al., 2018). On the lower axes, mean differences are plotted as bootstrap sampling distributions. Each mean difference is depicted as a dot. Each 95% confidence interval is indicated by the ends of the vertical error bars. Letters beneath each distribution refer to statistically indistinguishable groups after performing a Kruskall-Wallis test followed by a post hoc Mann-Whitney U-test (p<0.05).

### FACS

Fluorescence-activated cell sorting (FACS) was used to enrich eGFP labelled DANs from pupal/adult. The following genotypes were used for sorting: wild type *(THD’GAL4> UAS-eGFP), Orai^E180A^ (THD’GAL4> Orai^E180A^)*, Set2^IR-1^ *(THD’GAL4> Set2^IR-1^)*, *Orai^E180A^; Set2^OE^ (THD’GAL4> Orai^E180A^; Set2^OE^), Set2^OE^ (THD’GAL4> Set2^OE^), Trl^IR-1^ (THD’GAL4> Trl^IR-1^)*. Approximately 100 pupae per sample were washed in 1×PBS and 70% ethanol. Pupal/adult CNSs were dissected in Schneider’s medium (ThermoFisher Scientific) supplemented with 10% fetal bovine serum, 2% PenStrep, 0.02 mM insulin, 20 mM glutamine and 0.04 mg/ml glutathione. Post dissection, the CNSs were treated with an enzyme solution [0.75 g/l Collagenase and 0.4 g/l Dispase in Rinaldini’s solution (8 mg/ml NaCl, 0.2 mg/ml KCl, 0.05 mg/ml NaH2PO4, 1 mg/ml NaHCO3 and 0.1 mg/ml glucose)] at room temperature for 30 min. They were then washed and resuspended in ice-cold Schneider’s medium and gently triturated several times using a pipette tip to obtain a single-cell suspension. This suspension was then passed through a 40 mm mesh filter to remove clumps and kept on ice until sorting (less than 1 h). Flow cytometry was performed on a FACS Aria Fusion cell sorter (BD Biosciences) with a 100 mm nozzle at 60 psi. The threshold for GFP-positive cells was set using dissociated neurons from a non GFP-expressing wild-type strain (*Canton S*). The same gating parameters were used to sort other genotypes in the experiment. GFP-positive cells were collected directly in TRIzol and then frozen immediately in dry ice until further processing. Details for all reagents used are listed in Supplementary Table 1b.

### RNA-Seq

RNA from at least 600 sorted *THD’* DANs from the relevant genotypes, expressing *UAS-eGFP* was subjected to 14 cycles of PCR amplification (SMARTer Seq V4 Ultra Low Input RNA Kit; Takara Bio). 1 ng of amplified RNA was used to prepare cDNA libraries (Nextera XT DNA library preparation kit; Illumina). cDNA libraries for 4 biological replicates for both control (*THD’GAL4/+*) and experimental (*THD’GAL4>UAS-Orai^E180A^*) genotypes were run on a Hiseq2500 platform. 50-70 million unpaired sequencing reads per sample were aligned to the dm6 release of the *Drosophila* genome using HISAT2(Kim et al., 2015, 2019) and an overall alignment rate of 95.2-96.8% was obtained for all samples. Featurecounts(Liao et al., 2014) was used to assign the mapped sequence reads to the genome and obtain read counts. Differential expression analysis was performed using three independent methods: DESeq2(Love et al., 2014), limma-voom(Ritchie et al., 2015), and edgeR(Robinson et al., 2009). A fold change cutoff of a minimum twofold change was used. Significance cutoff was set at an FDR-corrected P value of 0.05 for DESeq2 and edgeR. Volcanoplots were generated using VolcaNoseR (https://huygens.science.uva.nl/). Comparison of gene lists and generation of Venn diagrams was performed using Whitehead BaRC public tools (http://jura.wi.mit.edu/bioc/tools/). Gene ontology analysis for molecular function was performed using DAVID(Huang et al., 2008, 2009). Developmental gene expression levels were measured for downregulated genes using FlyBase(Larkin et al., 2020) and DGET(Hu et al., 2017) (www.flyrnai.org/tools/dget/web/), and were plotted as a heatmap using ClustVis (Metsalu and Vilo, 2015) (https://biit.cs.ut.ee/clustvis/).

### qRT-PCRs

Central nervous systems (CNSs) of the appropriate genotype and age were dissected in 1× phosphate-buffered saline (PBS; 137 mM NaCl, 2.7 mM KCl, 10 mM Na_2_HPO_4_ and 1.8 mM KH_2_PO_4_) prepared in double distilled water treated with DEPC. Each sample consisted of five CNS homogenized in 500 µl of TRIzol (Ambion, ThermoFisher Scientific) per sample. At least three biological replicate samples were made for each genotype. After homogenization the sample was kept on ice and either processed further within 30 min or stored at −80°C for up to 4 weeks before processing. RNA was isolated following the manufacturer’s protocol. Purity of the isolated RNA was estimated by NanoDrop spectrophotometer (ThermoFisher Scientific) and integrity was checked by running it on a 1% Tris-EDTA agarose gel. Approximately 100 ng of total RNA was used per sample for cDNA synthesis. DNAse treatment and first-strand synthesis were performed as described previously(Pathak et al., 2015). Quantitative real time PCRs (qPCRs) were performed in a total volume of 10 µl with Kapa SYBR Fast qPCR kit (KAPA Biosystems) on an ABI 7500 fast machine operated with ABI 7500 software (Applied Biosystems). Technical duplicates were performed for each qPCR reaction. The fold change of gene expression in any experimental condition relative to wild type was calculated as 2−ΔΔCt, where ΔΔC_t_=[C_t_ (target gene) –C_t_ (rp49)] Expt. – [C_t_ (target gene) – C_t_ (rp49)]. Primers specific for rp49 and ac5c were used as internal controls. Sequences of all primers used are provided in Supplementary Table 1c.

### Functional imaging

Adult brains were dissected in adult hemolymph-like saline (AHL: 108 mM NaCl, 5 mM KCl, 2 mM CaCl2, 8.2 mM MgCl_2_, 4 mM NaHCO_3_, 1 mM NaH_2_PO_4_, 5 mM Trehalose, 10 mM Sucrose, 5 mM Tris, pH 7.5) after embedded in a drop of 0.1% low-melting agarose (Invitrogen). Embedded brains were bathed in AHL and then subjected to functional imaging. The genetically encoded calcium sensor *GCaMP6m* (Chen et al., 2013) was used to record changes in intracellular cytosolic Ca^2+^ in response to stimulation with carbachol or KCl. The GRAB-DA sensor (Sun et al., 2018) was used to measure evoked dopamine release. Images were taken as a time series on a xy plane at an interval of 1 s using a 20× objective with an NA of 0.7 on an Olympus FV3000 inverted confocal microscope. A 488 nm laser line was used to record GCaMP6m / GRAB-DA measurements. All live-imaging experiments were performed with at least 10 independent brain preparations. For measuring evoked responses, KCl/Carbachol, was added on top of the samples. For optogenetics experiments, flies were reared on medium supplemented with 200 μM ATR (all trans retinal), following which neuronal activation was achieved using a 633-nm laser line to stimulate *CsChrimson* (Klapoetke et al., 2014) while simultaneously acquiring images with the 488 nm laser line, and the images were acquired every 1 s. The raw images were extracted using FIJI (Schindelin et al., 2012) (based on ImageJ version 2.1.0/1.53c). ΔF/F was calculated from selected regions of interest (ROIs) using the formula ΔF/F=(Ft-F0)/F0, where Ft is the fluorescence at time t and F0 is baseline fluorescence corresponding to the average fluorescence over the first 40-time frames. Mean ΔF/F time-lapses were plotted using PlotTwist (Goedhart, 2020) (https://huygens.science.uva.nl/PlotTwist/). A shaded error bar around the mean indicates the 95% confidence interval for CCh (50 µM) or KCl (70 mM) responses. Peaks for individual cellular responses for each genotype was calculated from before or after the point of stimulation, using Microsoft Excel and plotted as a paired plot using SuperPlotsOfData (Goedhart, 2021) (https://huygens.science.uva.nl/SuperPlotsOfData/). Boxplots were plotted using PlotsOfData (Postma and Goedhart, 2019) (https://huygens.science.uva.nl/PlotsOfData/).

### Immunohistochemistry

Immunostaining of larval *Drosophila* brains was performed as described previously(Daniels et al., 2008). Briefly, adult brains were dissected in 1x phosphate buffered saline (PBS) and fixed with 4% Paraformaldehyde. The dissected brains were washed 3-4 times with 0.2% phosphate buffer, pH 7.2 containing 0.2% Triton-X 100 (PTX) and blocked with 0.2% PTX containing 5% normal goat serum (NGS) for 2 hours at room temperature. Respective primary antibodies were incubated overnight (14–16 hr) at 4°C. After washing 3-4 times with 0.2% PTX at room temperature, they were incubated in the respective secondary antibodies for 2 hr at room temperature. The following primary antibodies were used: chick anti-GFP antibody (1:10,000; A6455, Life Technologies), mouse anti-nc82 (anti-brp) antibody (1:50), rabbit anti-H3K36me3, mouse anti-RFP. Secondary antibodies were used at a dilution of 1:400 as follows: anti-rabbit Alexa Fluor 488 (#A11008, Life Technologies), anti-mouse Alexa Fluor 488 (#A11001, Life Technologies). Confocal images were obtained on the Olympus Confocal FV1000 microscope (Olympus) with a 40x, 1.3 NA objective or with a 60x, 1.4 NA objective. Imaging for axonal arbors within the mushroom body γ lobe was performed on the Zeiss LSM980 system with AiryScan 2. Images were visualized using either the FV10-ASW 4.0 viewer (Olympus) or Fiji (Schindelin et al., 2012). Details for all reagents used are listed in Supplementary Table 1b.

### Western Blotting

Adult CNSs of appropriate genotypes were dissected in ice-cold PBS. Between 5 and 10 brains were homogenized in 50 μl of NETN buffer [100 mM NaCl, 20 mM Tris-Cl (pH 8.0), 0.5 mM EDTA, 0.5% Triton-X-100, 1× Protease inhibitor cocktail (Roche)]. The homogenate (10-15 μl) was run on an 8% SDS-polyacrylamide gel. The protein was transferred to a PVDF membrane by standard semi-dry transfer protocols (10 V for 10 min). The membrane was incubated in the primary antibody overnight at 4°C. Primary antibodies were used at the following dilutions: rabbit anti-H3K36me3 (1:5000, Abcam, ab9050) and rabbit anti-H3 (1:5000, Abcam, ab12079). Secondary antibodies conjugated with horseradish peroxidase were used at dilution of 1:3000 (anti-rabbit HRP; 32260, Thermo Scientific). Protein was detected by a chemiluminescent reaction (WesternBright ECL, Advansta K12045-20). Blots were first probed for H3K36me3, stripped with a solution of 3% glacial acetic acid for 10 min, followed by re-probing with the anti-H3 antibody.

### In silico ChIP-Seq Analysis

H3K36me3 enrichment data was obtained from a ChIP-chip dataset (ID_301) generated in *Drosophila* ML-DmBG3-c2 cells submitted to modEncode(Celniker et al., 2009; modENCODE Consortium et al., 2010). Enrichment scores for genomic regions were calculated using ‘computematrix’ and plotted as a tag density plot using ‘plotHeatmap’ from deeptools2(Ramírez et al., 2016). All genes were scaled to 2 kb with a flanking region of 250 bp on either end. A 50 bp length of non-overlapping bins was used for averaging the score over each region length. Genes were sorted based on mean enrichment scores and displayed on the heatmap in descending order.

### Statistical tests

Non-parametric tests were employed to test significance for data that did not follow a normal distribution. Significant differences between experimental genotypes and relevant controls were tested either with the Kruskal-Wallis test followed by Dunn’s multiple comparison test (for multiple comparisons) or with Mann-Whitney U tests (for pairwise comparisons). Data with normal distribution were tested by the Student’s T-test. Statistical tests and p-values are mentioned in each figure legend. For calculation and representation of effect size, estimationstats.com was used.

## Acknowledgements

We acknowledge the Central Imaging and Flow Cytometry Facility (CIFF, NCBS) for maintenance of microscopes and the *Drosophila* facility (Flyfacility, NCBS) for Fly stock maintenance and development of transgenics. We thank Nandashree KS for helping with super resolution imaging on the Zeiss 980 Airyscan system.

## Data availability

The RNA Sequencing data has been submitted to GEO under accession number GSE230134. All materials including source data have been made available.

