## Supplementary material for "Orai mediated Calcium entry determines activity of central dopaminergic neurons by regulation of gene expression": Figure supplements

Figure 1- Figure Supplement 1

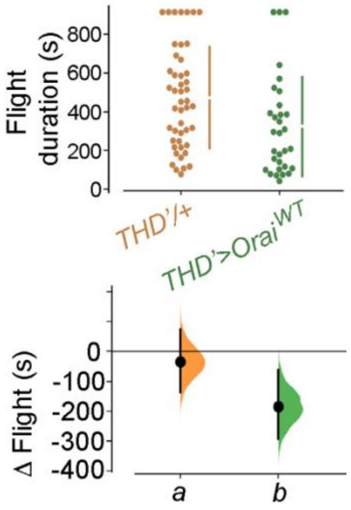

Figure 1- Figure Supplement 2

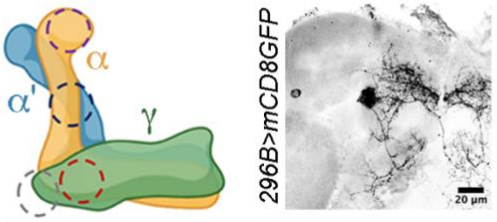


| | MB504B | MB296B | MB438B |
|---|---|---|---|
| γ1p | ● | | ● |
| γ2α'1 | ● | ● | |
| α'2α2 | ● | | ● |
| α3 | ● | | ● |

Figure 1- Figure Supplement 3

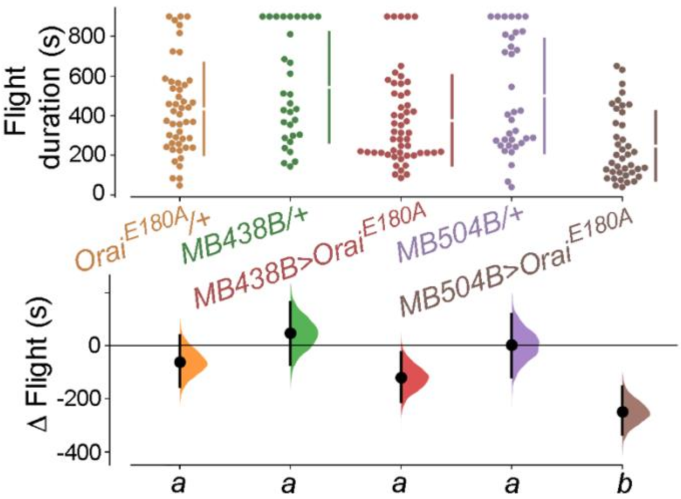

Figure 1- Figure Supplement 4

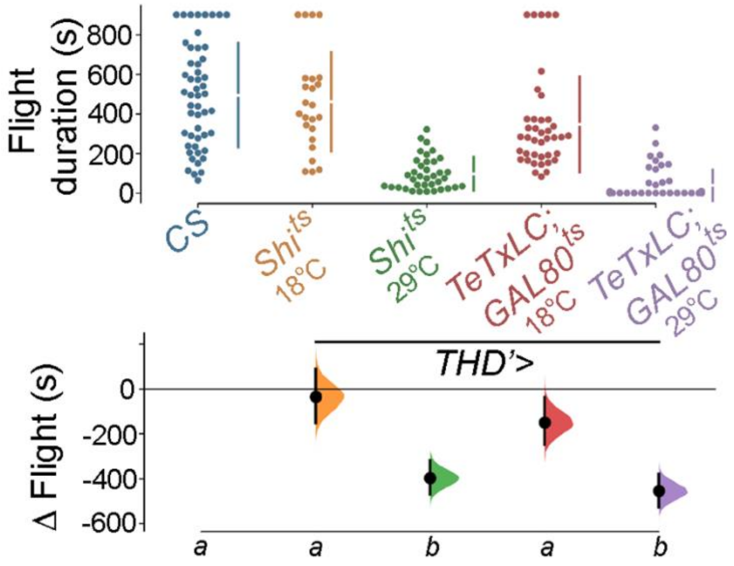

Figure 1- Figure Supplement 5

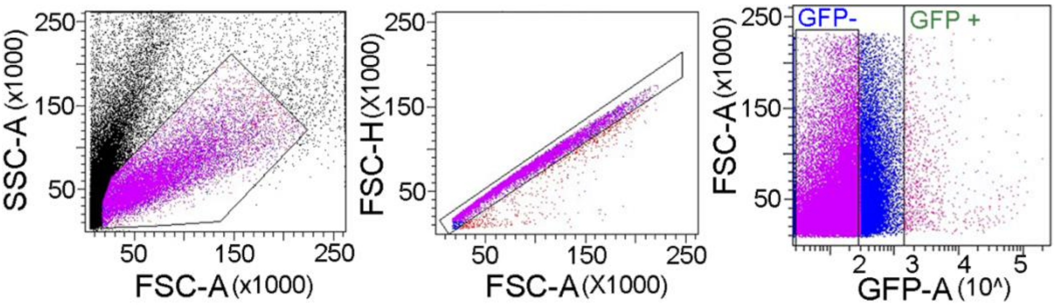

Figure 1- Figure Supplement 6

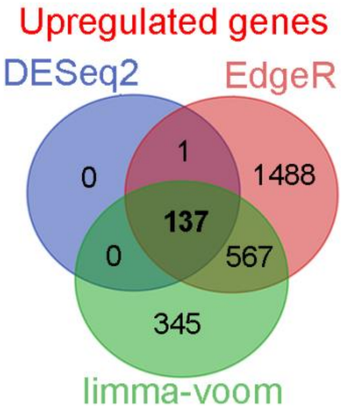

Figure 1- Figure Supplement 7

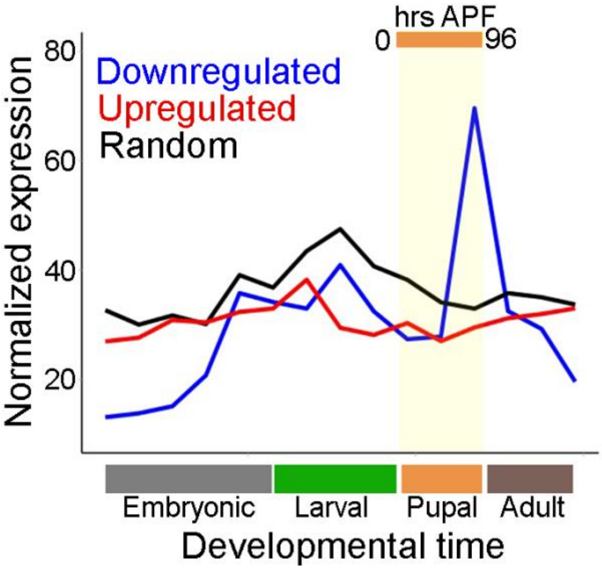

Figure 1- Figure Supplement 8

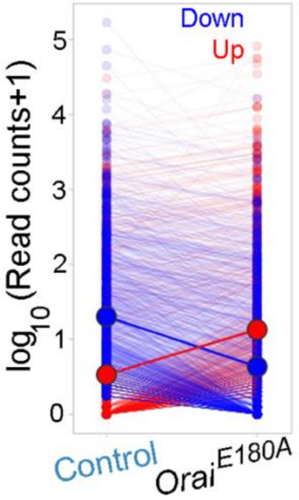

Figure 1- Figure Supplement 9

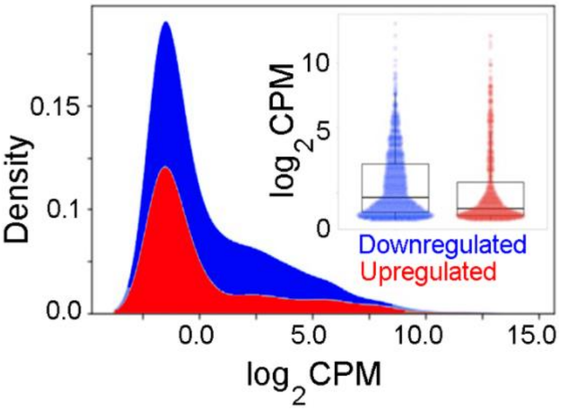

Figure 2- Figure Supplement 1

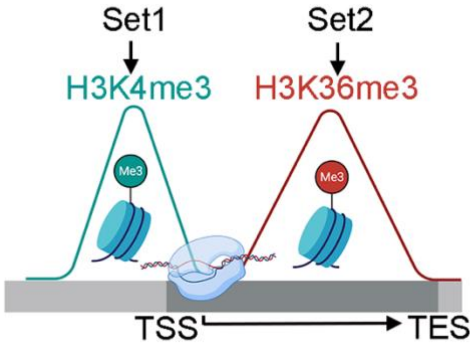

Figure 2- Figure Supplement 2

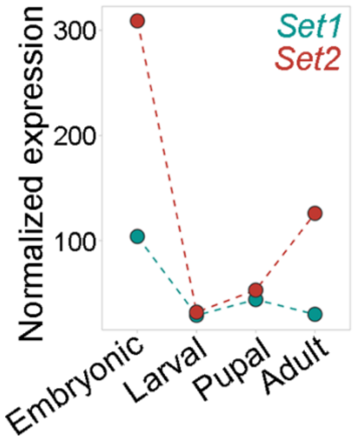

Figure 2- Figure Supplement 3

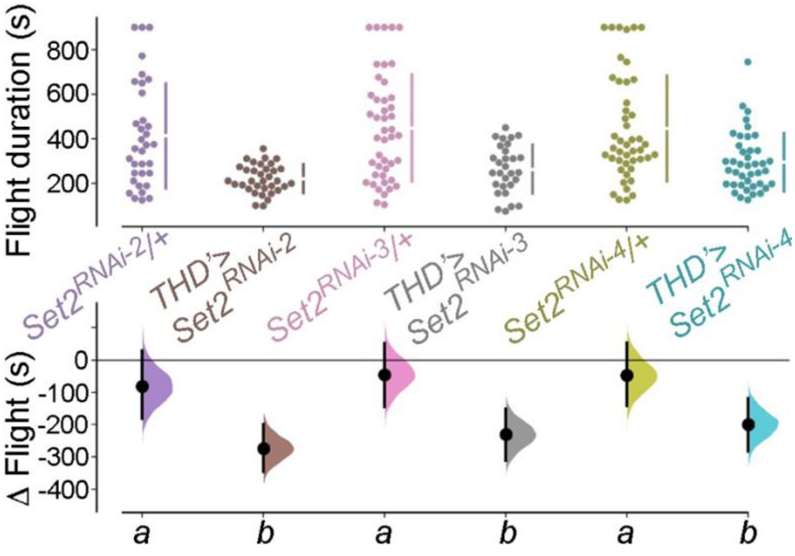

Figure 2- Figure Supplement 4

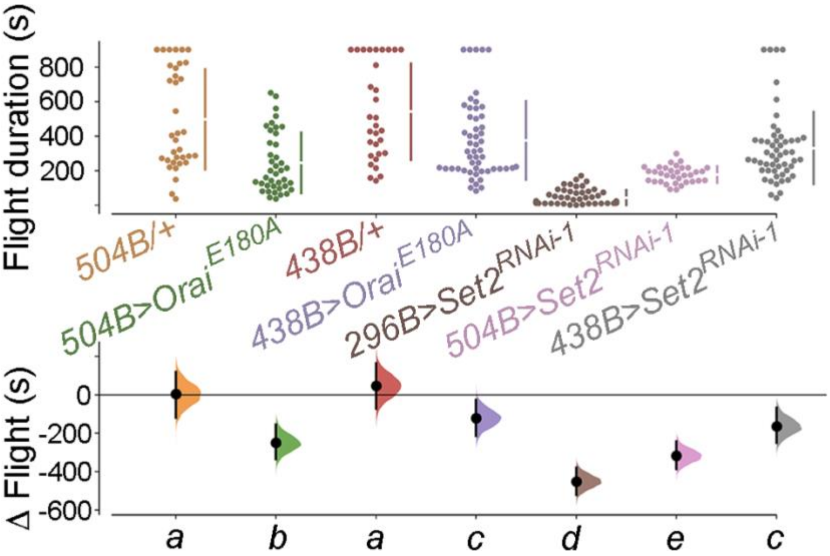

Figure 2- Figure Supplement 5

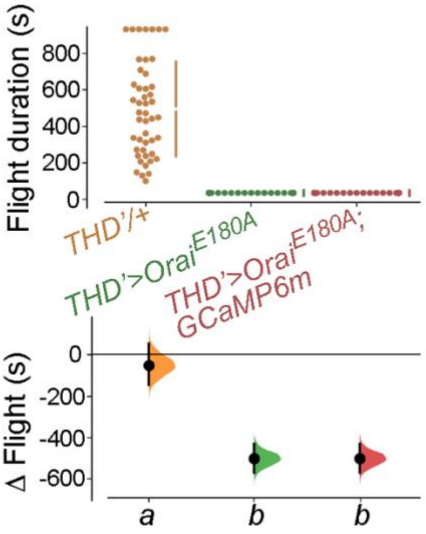

Figure 2- Figure Supplement 6

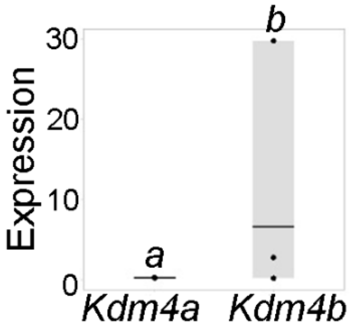

Figure 2- Figure Supplement 7

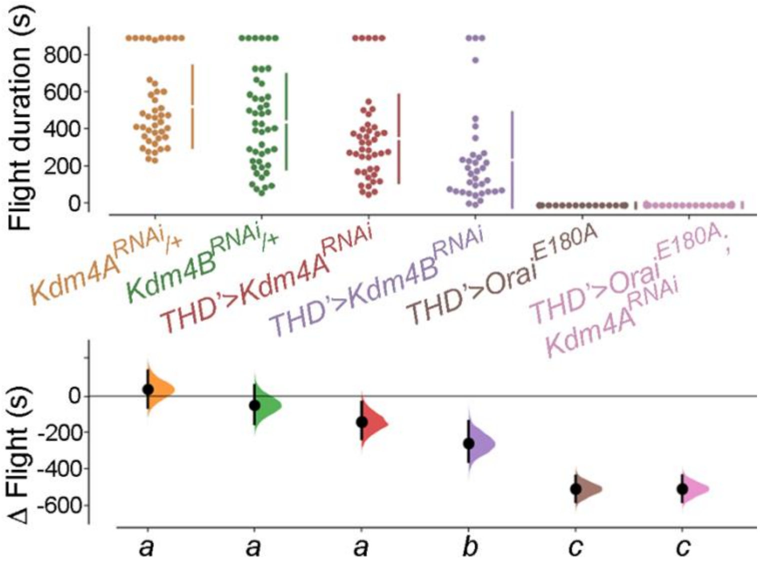

Figure 2- Figure Supplement 8

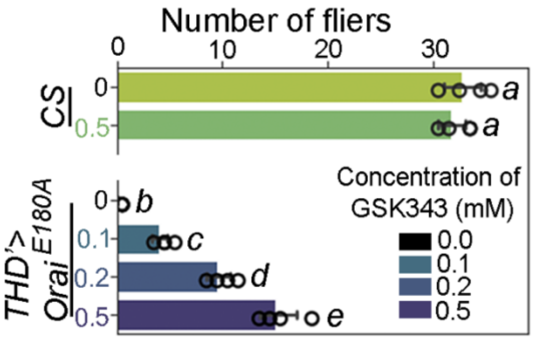

Figure 3- Figure Supplement 1

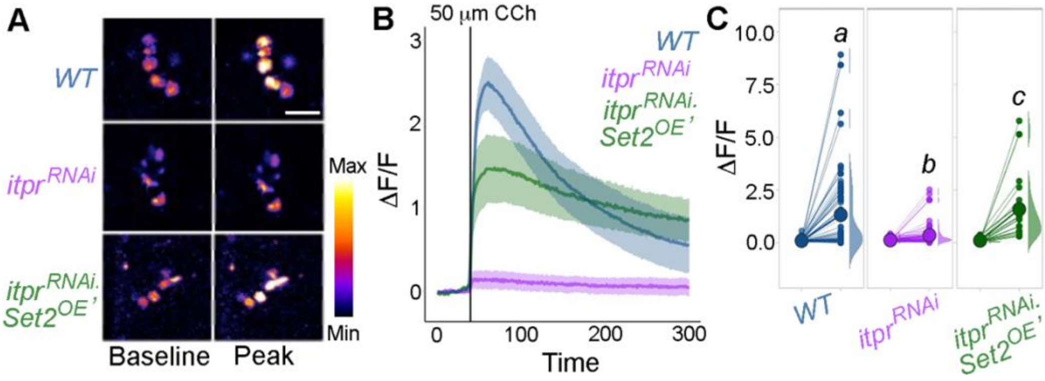

Figure 4- Figure Supplement 1

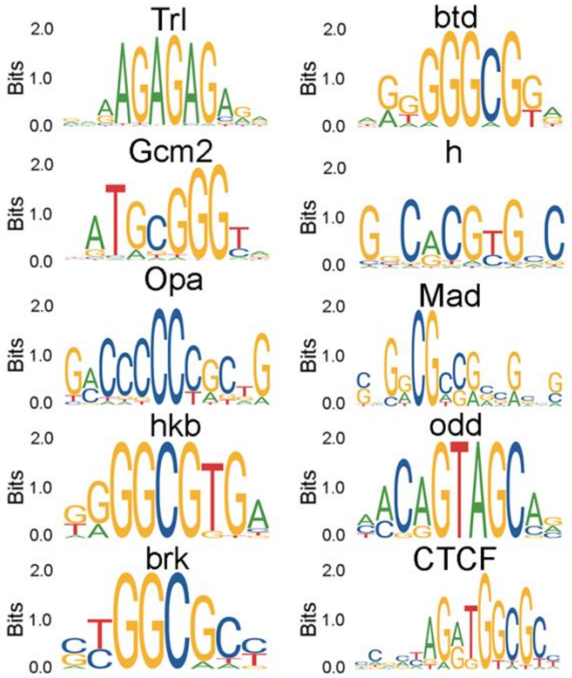

Figure 4- Figure Supplement 2

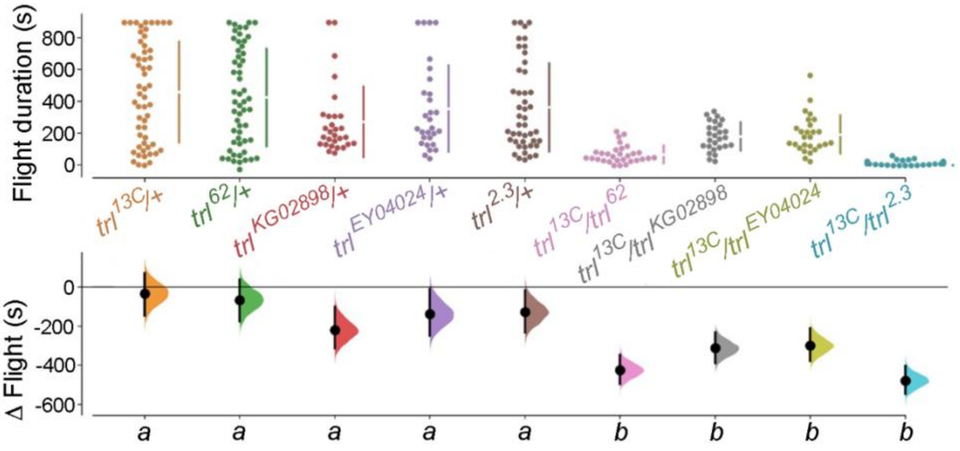

Figure 4- Figure Supplement 3

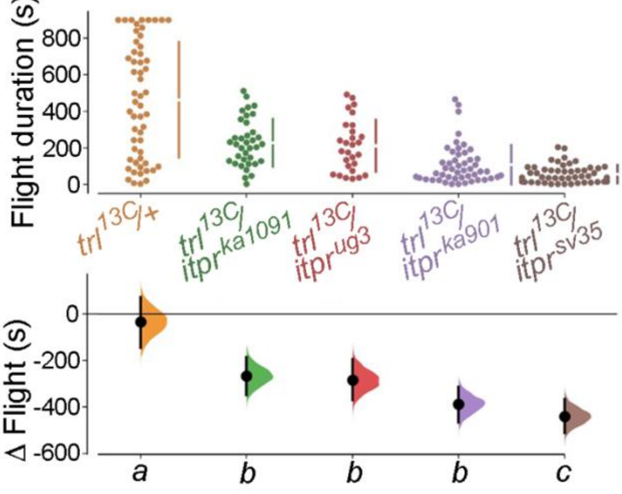

Figure 4- Figure Supplement 4

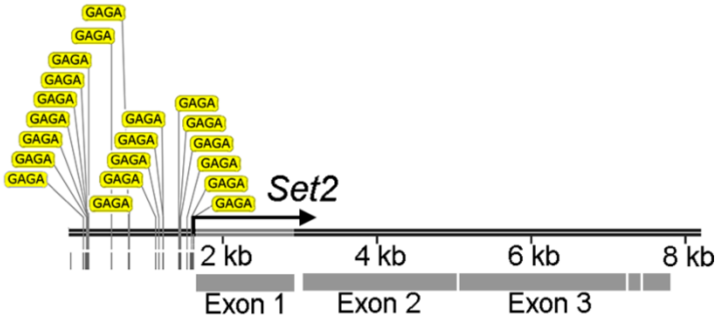

Figure 5- Figure Supplement 1

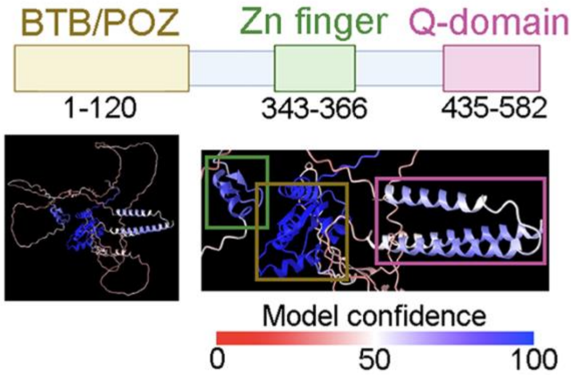

Figure 5- Figure Supplement 2

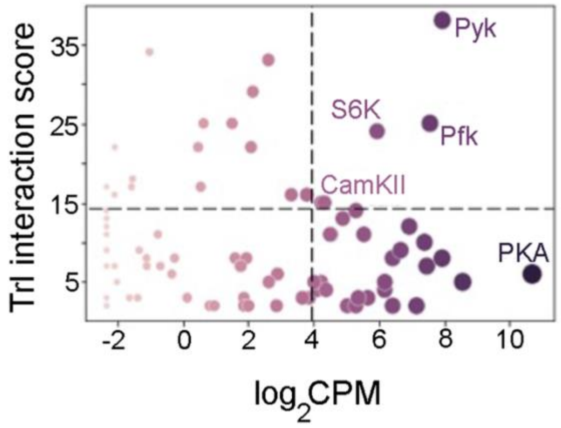

Figure 5- Figure Supplement 3

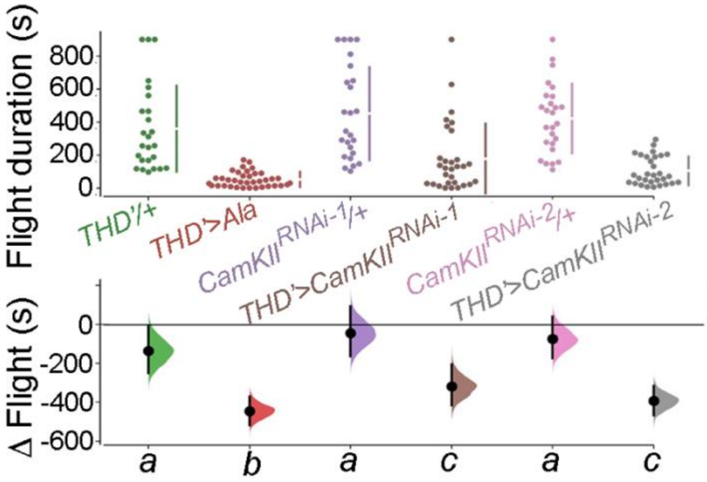

Figure 5- Figure Supplement 4

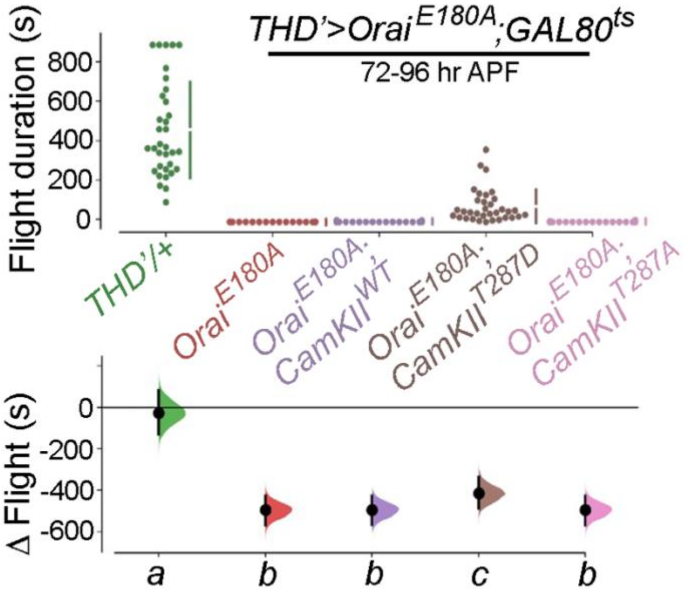

Figure 7- Figure Supplement 1

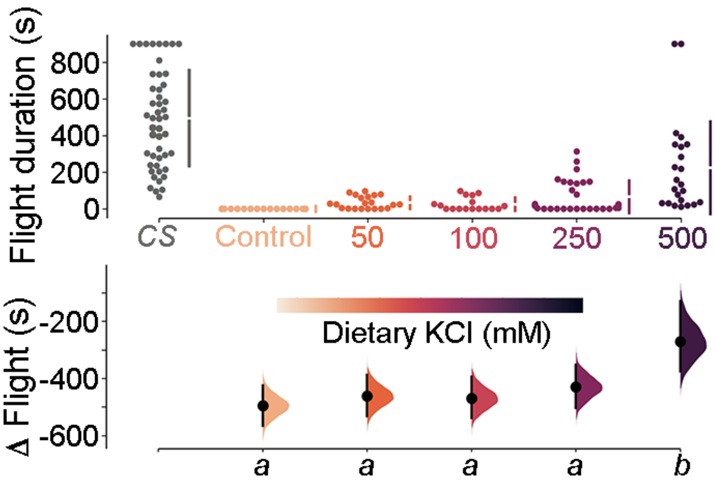

Figure 7- Figure Supplement 2

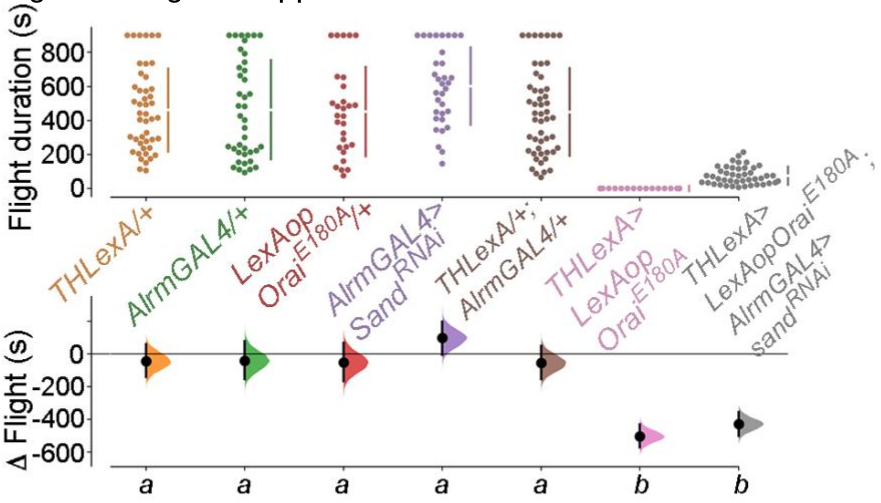

Figure 7- Figure Supplement 3

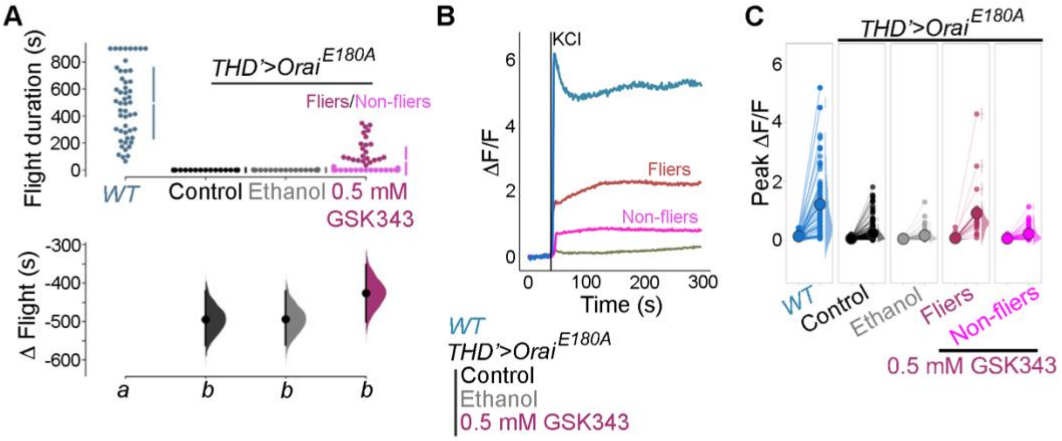
