## Supplementary File 1 for "Orai mediated Calcium entry determines activity of central dopaminergic neurons by regulation of gene expression"

**Supplementary File 1a: Fly Stocks**

| **Stocks** | **Description** | **Reference** |
| --- | --- | --- |
| **WT** | | |
| *Canton S* |  |  |
| **UAS lines** | | |
| *UASitpr^IR^* | RNAi line for *itpr* | 1063-R2 from NIG |
| *UASitpr^IR^; UASDicer* | RNAi line for *itpr* combined with dicer | 1063-R2 from NIG; BL24648 |
| *UASdStim ^IR^; UASdicer* | RNAi line for *dStim* combined with dicer | VDRC: 47073 |
| *UAS Set2 ^RNAi-1^* | Set2 RNAi | BDSC: 24108 |
| *UAS Set2 ^RNAi-2^* | Set2 RNAi | BDSC: 55221 |
| *UAS Set2 ^RNAi-3^* | Set2 RNAi | BDSC: 33706 |
| *UAS Set2 ^RNAi-4^* | Set2 RNAi | BDSC: 42511 |
| *UAS Trl ^RNAi-1^* | Trl RNAi | BDSC: 40940 |
| *UAS Trl ^RNAi-2^* | Trl RNAi | BDSC: 67265 |
| *UAS cac ^RNAi^* | *Cac* RNAi | BDSC:77174 |
| *UAS ca-α1D ^RNAi^* | *ca-α1D* RNAi | BDSC:25830 |
| *UAS ca-α1T ^RNAi^* | *ca-α1T* RNAi | BDSC:26251 |
| *UAS ca-β ^RNAi^* | *ca-β* RNAi | BDSC:29575 |
| *UAS-CaMKII* | WT *CamKII* overexpression | BDSC: 29662 |
| *UAS-CaMKII-T287D* | *CamKII* dominant-active | BDSC: 29665 |
| *UAS-CaMKII-T287A* | *CamKII* dominant-active control | BDSC: 29664 |
| *UAS-CaMKII^RNAI-1^* | *CamKII* RNAi | BDSC: 35330 |
| *UAS-CaMKII^RNAI-2^* | *CamKII* RNAi | VDRC: 38930 |
| *UAS Kdm4A ^RNAi^* | *Kdm4A* RNAi | BDSC:29376 |
| *UAS Kdm4B ^RNAi^* | *Kdm4B* | BDSC:34629 |
| *UAS-Ala* | *CamKII* inhibitory peptide | BDSC: 29666 |
| *UAS Set2* | *Set2* overexpression | Mitra et al., 2021 |
| *UAS H2BmRFP* | Nuclear tagged RFP | Gomes et al., 2009 |
| *UAS mCD8GFP* | Membrane tagged GFP | BDSC:79626 |
| *UASeGFP* | Cytosolic GFP | Gift from Michael Rosbash |
| *UAS GCaMP6m* | Genetically encoded calcium indicator | BDSC: BL42748 |
| *UAS Stim* | Stim overexpression | Agrawal et al., 2009 |
| *UAS Shibire^ts^* | Temperature sensitive shibire | BDSC:30012 |
| *UAS TNT* | TetXLC for inhibiting synaptic transmission | BDSC:28837 |
| *UAS CsChrimson* | Red light sensitive channel for optogenetic activation | BDSC:55135 |
| *UAS GtACR2* | Green light sensitive channel for optogenetic inhibition | BDSC:92984 |
| *UAS TrpA1* | Temperature sensitive TrpA1 for neuronal activation | Kang et al., 2012 |
| *UAS Kir2.1* | Kir2.1 expression for neuronal inhibition | BDSC:91802 |
| *UAS-NachBac* | Bacterial Sodium channel for neuronal activation | BDSC:9468 |
| *UAS cac* | *Cacophony* overexpression | BDSC:8581 |
| **GAL4 lines** |  |  |
| *elav^C155^GAL4* | Pan-neuronal driver | Luo et al., 1994 |
| *tubulinGAL80^ts^/tubulinGAL80^ts^; tubulinGAL80^ts^/tubulinGAL80^ts^* | Temperature sensitive GAL80 under the tubulin promoter | McGuire et al., 2003 |
| *nsybGAL4* | Pan neuronal *GAL4* | BDSC: 5136 |
| *THD’GAL4* | *GAL4* marking all dopaminergic neurons | Liu et al., 2012 |
| *MB296BGAL4* | Split- *GAL4* marking PPL1- γ2α′1 dopaminergic neurons | BDSC: 63308 |
| *MB504BGAL4* | Split- *GAL4* marking PPL1- α′2α2 γ2α′1α3 γ1 pedc dopaminergic neurons | BDSC: 68329 |
| *MB438BGAL4* | Split- *GAL4* marking PPL1- α′2α2α3γ1 pedc dopaminergic neurons | BDSC: 68326 |
| **Mutants** |  |  |
| *itpr^ug3^* | Hypomorphic mutant allele for *itpr* | Joshi et al., 2004 |
| *itpr^ka1091^* | Hypomorphic mutant allele for *itpr* | Joshi et al., 2004 |
| *itpr^ka901^* | Hypomorphic mutant allele for *itpr* | Joshi et al., 2004 |
| *itpr^sv35^* | Hypomorphic mutant allele for *itpr* | Joshi et al., 2004 |
| *trl^13C^* | Hypomorphic mutant allele for *Trl* | Farkas et al, 1994 |
| *trl^62^* | Hypomorphic mutant allele for *Trl* | Farkas et al, 1994 |
| *trl^KG02898^* | Hypomorphic mutant allele for *Trl* | Bellen et al., 2004 |
| *trl^EY04024^* | Hypomorphic mutant allele for *Trl* | Rorth et al., 1996; Quijano et al., 2016 |

**Supplementary File 1b: Reagents**

| **Reagent** | **Designation** | **Source** | **Identifier** |
| --- | --- | --- | --- |
| Antibody | Rabbit anti H3K36me3 (polyclonal) | Abcam | RRID:AB_306966, Cat# ab9050 |
| Antibody | Rabbit anti H3 (polyclonal) | Abcam | RRID:AB_302613, Cat# ab1791 |
| Antibody | Rabbit anti GFP (polyclonal) | Life Technologies, Thermo Fisher | RRID:AB_221570,Cat # A-6455 |
| Antibody | Mouse anti RFP (polyclonal) | Rockland | RRID:AB_2209751, Cat# 200-301-379 |
| Antibody | Mouse anti-bruchpilot (monoclonal) | Gift from Eric Buchner, University of Wuerzburg | |
| Antibody | Anti-mouse Alexa Fluor 568 (polyclonal) | Life Technologies, ThermoFisher Scientific | Cat# A-11004, RRID:AB_2534072 |
| Antibody | Anti-rabbit Alexa Fluor 488 (polyclonal) | Life Technologies, ThermoFisher Scientific | Cat# A-11008, RRID:AB_143165 |
| Antibody | Anti-rabbit HRP (polyclonal) | ThermoFisher Scientific | Cat#32260, RRID:AB_1965959 |
| Chemical compound | Low melt Agar | Invitrogen | Cat# 16520–050 |
| Chemical compound | Carbachol | Sigma Aldrich | Cat# C4382 |
| Chemical compound | All trans Retinal | Sigma Aldrich | Cat# R2500 |
| Chemical compound | TTX | Sigma Aldrich | Cat #582231 |
| Chemical compound | Nimodipine | Sigma Aldrich | Cat #N149 |
| Chemical compound | GSK343 | Cell Signallng Technology | Cat #66244 |

**Supplementary File 1c: Primer Sequences**

| **Primer** | **Sequence (5’-3’)** |
| --- | --- |
| rp49-F | CGGATCGATATGCTAAGCTGT |
| rp49-R | GCGCTTGTTCGATCCGTA |
| dStim-F | GAAGCAATGGATGTGGTTCTG |
| dStim-R | CCGAGTTCGATGAACTGAGAG |
| Set2-F | CCAATGCCACCGAGTGTTAC |
| Set2-R | TCCTTGCGATACGGACGC |
| itpr-F | CCAGGGTTTGCGAAATGGC |
| itpr-R | CAGGTCGTCTTCAGAATGGC |
| mAchR-F | ATGACACCTGGCGACGTCC |
| mAchr-R | CGCAATGCACCACTCCTTG |
| GFP-F | TGGCCCTGTCCTTTTACCAG |
| GFP-R | CCATGCCATGTGTAATCCCAG |
| Orai-F | GAGATAGCCATCCTGTGCTGG |
| Orai-R | CGGATGCCCGAGACTGTC |
| tub-F | CCAAGGGTCATTACACAGAGG |
| tub-R | ATCAGCAGGGTTCCCATACC |
| cac-F | TGTTCGATTGCGTCGTGAAC |
| cac-R | TGGCACTCTGCGGAAGTATG |
| ca-alpha1D-F | GCGAATGCCATTAACTATGACAAC |
| ca-alpha1D-R | ACTCGGAGTCGCAGTATTTACC |
| ca-alpha1T-F | GCGGTGAACTGAAAAGAGAAC |
| ca-alpha1T-R | CAAATCGGGTGGTTTACGATGG |
| ca-beta-F | GGAAGCCAAGATACCCGAGC |
| ca-beta-R | CGAAGATCTCGGCGAAGAGG |
| hk-F | CTGTTCGACATCTCGGAGGC |
| hk-R | GGTGCTCCAGTAGACCTTCG |
| Slo-F | GCTGGACCTGAACACATGGA |
| Slo-R | TGCGGCTCCTACTTATCTGC |
| eag-F | AGGATGTTTCCCTGCTCGTG |
| eag-R | TGGGTCACTAACAACCTCGC |
| NaCP60E-F | GACTTTTCCGAGCTACGGGG |
| NaCP60E-R | TTAAGACAGCCGTCCTCACG |
